# Single cell transcriptomics of the developing zebrafish lens and identification of putative controllers of lens development

**DOI:** 10.1101/2020.12.09.418533

**Authors:** Dylan Farnsworth, Mason Posner, Adam Miller

## Abstract

The vertebrate lens is a valuable model system for investigating the gene expression changes that coordinate tissue differentiation due to its inclusion of two spatially separated cell types, the outer epithelial cells and the deeper denucleated fiber cells that they support. Zebrafish are a useful model system for studying lens development given the organ’s rapid development in the first several days of life in an accessible, transparent embryo. While we have strong foundational knowledge of the diverse lens crystallin proteins and the basic gene regulatory networks controlling lens development, no study has detailed gene expression in a vertebrate lens at single cell resolution. Here we report an atlas of lens gene expression in zebrafish embryos at single cell resolution through five days of development, identifying a number of novel regulators of lens development as potential targets for future functional studies. Our temporospatial expression data address open questions about the function of α-crystallins during lens development and provides the first detailed view of β- and γ-crystallin expression in and outside the lens. We describe subfunctionalization in transcription factor genes that occur as paralog pairs in the zebrafish. Finally, we examine the expression dynamics of cytoskeletal, RNA-binding, and transcription factors genes, identifying a number of novel patterns. Overall these data provide a foundation for identifying and characterizing lens developmental regulatory mechanisms and revealing targets for future functional studies with potential therapeutic impact.

## 1. Introduction

The vertebrate ocular lens is a valuable model system for studying development, physiology, disease and gene evolution. The presence of two terminal cell types, an outer single-layered epithelium and inner fiber cell mass, simplifies the examination of signaling processes leading to tissue differentiation (Grainger, 1992). The ease with which the lens can be imaged, both intact and histologically, makes it an excellent tool for identifying anatomical abnormalities and imaging cellular structure. Additionally, the co-option of numerous non-eye related genes during the early evolution of the vertebrate lens provides a unique opportunity to study the evolution of development and of proteins for diverse roles (Wistow and Piatigorsky, 1988). Most notably, small heat shock proteins became the α-crystallins of the lens, serving both as structural refractive elements while also inhibiting protein aggregation that would otherwise lead to lens cataract, the leading cause of human blindness worldwide.

Zebrafish have been an important species for studying eye development and function (Bibliowicz et al., 2011; Morris, 2011). The evolutionary conservation of gene expression and protein content between the zebrafish and mammalian lens facilitates the use of zebrafish as a model system for vision research (Greiling et al., 2009; Posner et al., 2008; Richardson et al., 2016). Some differences in eye development between fish and mammals, such as the mechanism of lens delamination from the surface ectoderm and the presence of duplicated gene paralogs, allows for comparative studies (Clemens et al., 2013; Greiling et al., 2010). Zebrafish are also an excellent model for mutational analysis due to their abundant and accessible embryos and availability of tools for gene editing (J. Liu et al., 2016). Gene targeting techniques have been used in zebrafish to interrogate the function of lens-associated crystallins (Mishra et al., 2018; Posner et al., 2013; Zou et al., 2015), aquaporins (Clemens et al., 2013; Froger et al., 2010; Vorontsova et al., 2018), transcription factors (Krall and Lachke, 2018; Shi et al., 2005; 2006), DNA methylating enzymes (Tittle et al., 2011), cytoskeletal proteins and most recently RNA-binding proteins (Shao et al., 2020).

Studies with mice have contributed to a strong foundational understanding of the molecular mechanisms underlying lens development. For example, mouse microarray gene expression data was used to develop the iSyTE database of genes with eye preferred expression, leading to the discovery of several novel lens-related transcription factors and the first known RNA-binding protein regulating lens development, TDRD7 (Agrawal et al., 2015; Audette et al., 2016; Kakrana et al., 2017; Lachke et al., 2011). RNA-Seq and proteomic analysis of mouse lenses microdissected to separate the outer, single layer epithelium from the inner elongated fiber cells provided detailed analyses of changes in gene expression through four developmental time points and identified multiple genes with increased expression in one of these two lens tissues (Zhao et al., 2018a; 2018b). In total, these studies validate an approach of using lens preferred gene expression as a tool for identifying potential novel developmental regulators that can then be tested by mutational analysis.

While these past studies in mouse, zebrafish and other vertebrate species have provided important insights into lens biology and development, current techniques that measure gene expression in individual cells open up new ways to visualize tissue preferred expression and dynamic changes in gene expression during tissue differentiation. No study to date has analysed lens development at single cell resolution. We recently published a single cell RNA-Seq (scRNA-Seq) atlas for zebrafish development, with over 44,000 cells from embryos at 1, 2 and 5 days post fertilization in 220 clusters annotated for specific tissue types (Farnsworth et al., 2019). Two of these clusters were identified as cells of the ocular lens. Here, we use this scRNA-Seq dataset to analyze changes in gene expression through these five days of zebrafish development. These embryonic stages span the time when the peripheral lens epithelium transitions to a simple cuboidal layer that both supports deeper fiber cells while also generating newly differentiating fiber cells at its posterior margin. We describe patterns of gene expression in the numerous and diverse crystallins, cytoskeletal and membrane proteins that contribute to lens transparency and the refractive power required for vision. We also compare the expression of known lens development regulatory genes between zebrafish and mammals and examine the impacts of duplicated zebrafish paralogs on regulatory networks. Our identification of genes with preferred expression in zebrafish lens epithelium and fiber cells provides a list of possible new targets for investigation of lens development, differentiation, and function.

## 2. Materials and methods

### 2.1. Fish husbandry

Fish were maintained by the University of Oregon Zebrafish facility using standard husbandry techniques (Westerfield, 2007) under an Institutional Animal Care and Use Committee (IACUC) approved protocol AUP-18-31. Zebrafish, *Danio rerio*, were bred and maintained at 28°C on a 14 hr on and 10 hr off light cycle. Animal care was provided by the University of Oregon fish facility staff, and veterinary care was provided by Dr. Kathy Snell, DVM. Zebrafish are not yet sexually determined at these stages of development (Wilson et al., 2014). Embryos were collected from natural matings, staged and pooled (n = 15 per replicate). Animals used in this study were: *Tg(olig2:GFP)vu12* for 1 dpf, n = 2; 2 dpf, n = 1; 5 dpf, n = 2 samples and *Tg(elavl3:GCaMP6s*) for 2 dpf, n = 1 sample.

### 2.2. Embryo dissociation

Collagenase P was prepared to a 100 mg/mL stock solution in HBSS. Chemical dissociation was performed using 0.25% Trypsin, Collagenase P (2 mg/mL), 1 mM EDTA (pH 8.0), and PBS for 15 min at 28C with gently pipetting every 5 min. Dissociation was quenched using 5% calf serum, 1 mM CaCl2, and PBS. Cells were washed and resuspended in chilled (4C), 1% calf serum, 0.8 mM CaCl2, 50 U/mL penicillin, 0.05 mg/mL streptomycin, and DMEM and passed through a 40 μM cell strainer (Falcon) and diluted into PBS + 0.04% BSA to reduce clumping. A final sample cell concentration of 2000 cells per microliter, as determined on a Biorad TC20 cell counter, was prepared in PBS +0.04% BSA for cDNA library preparation.

### 2.3. Single-cell cDNA library preparation

Sample preparation was performed by the University of Oregon Genomics and Cell Characterization core facility (https://gc3f.uoregon.edu/). Dissociated cells were run on a 10X Chromium platform using 10x v.2 chemistry targeting 10,000 cells. The resulting cDNA libraries were amplified with 15 cycles of PCR and sequenced on either an Illumina HiSeq (5/6 samples) or an Illumina NextSeq (n = 1, 48 h dpf sample).

### 2.4. Computational analysis

The resulting sequencing data were analyzed using the 10X Cellranger pipeline, version 2.2.0 (Zheng et al., 2019) and the Seurat (Satija et al., 2015) software package for R, v3.4.4 (R Core Team, n.d.) using standard quality control, normalization, and analysis steps. Briefly, we aligned reads to the zebrafish genome, GRCz11_93, and counted expression of protein coding reads. The resulting matrices were read into Seurat where we performed PCA using 178 PCs. UMAP analysis was performed on the resulting dataset with 178 dimensions and a resolution of 18.0. For each gene, expression levels are normalized by the total expression, multiplied by a scale factor (10,000) and log-transformed. Differential gene expression analysis was performed using the FindMarkers function in Seurat using Wilcoxon rank sum test.

The lens cell clusters in the atlas were identified by mapping the expression of well-characterized lens crystallin and transcription factor genes. Fiber cell preferred β-crystallins and integral membrane proteins were used to identify that cell type while a known lens epithelial specific transcription factor, *foxe3*, was used to identify two epithelial cell clusters.

### 2.5. Analysis of gene expression shift in epithelial cells

We used gene ontology term (GO) analysis to compare the function of preferentially expressed genes between 1 and 2/5 dpf epithelial cells. The single cell atlas data were used to generate lists of genes with expression in our two identified epithelial cell clusters. Genes expressed in each cluster with adjusted p-values ≤ 0.05 were submitted to GOrilla to generate GO terms using the *Process* option (Eden et al., 2007; 2009).

## 3. Results

### 3.1. Well-characterized marker genes identified distinctly clustered lens epithelial and fiber cells

The 10X Genomics platform was used to identify and quantify mRNAs from 44,102 cells isolated from 1, 2 and 5 days post fertilization zebrafish embryos. Annotation of 220 cell clusters produced by UMAP non-linear dimensional reduction previously identified a lens cell region based on expression of lens specific crystallin protein genes (Farnsworth et al., 2019). Our first goal was to determine whether we could identify specific clusters for lens epithelial and fiber cells based on expression of known marker genes. For example, three genes known to be lens fiber cell enriched in mammals are the β-crystallin *cryba1a*, lactose binding *grifin*, and the aquaporin *mipa* (Hall and Mathias, 2014; Ogden et al., 1998; Wistow and Piatigorsky, 1988). These three genes showed shared expression in a specific set of cells that we identify as lens fiber cells (Fig 1A-D). The transcription factor *foxe3* is known to be specific to lens epithelial cells (Landgren et al., 2008; Shi et al., 2006), and in our single cell data it was expressed in two cell clusters (Fig. 1A, E), suggesting a difference in global gene expression among lens epithelial cells. These two separate epithelial clusters contained either 1 day post fertilization (dpf) or combined 2 and 5 dpf cells (Fig. 1F and G). There were no apparent differences in clustering between fiber cells at 1, 2 and 5 dpf (Fig. 1F and G).

**Figure 1.**
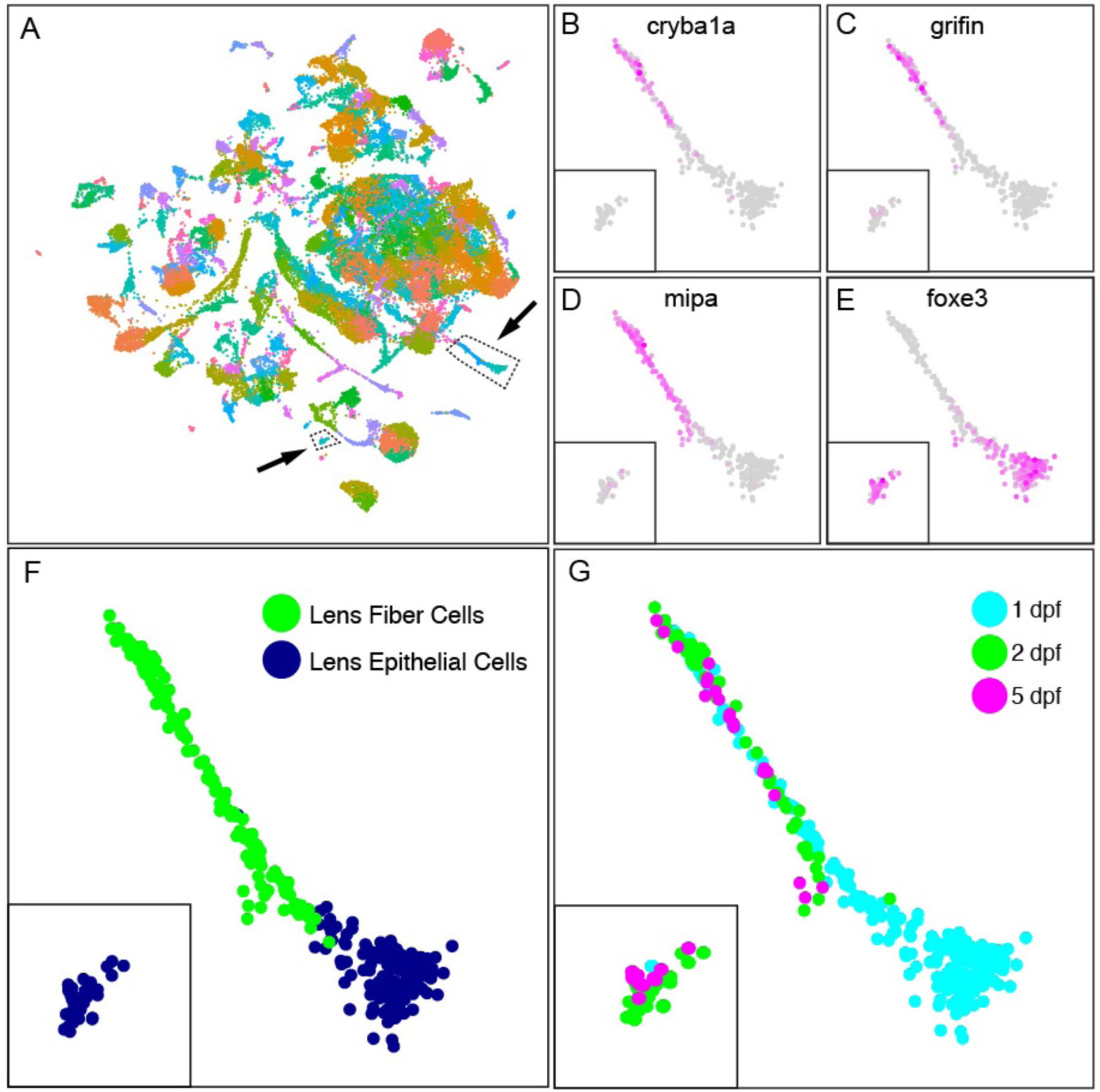
Expression of known lens marker genes identified lens epithelial and fiber cell clusters. Global single cell gene expression in 1, 2 and 5 dpf embryos was analyzed with UMAP to cluster cells with similar expression patterns (A). Two areas in the total atlas (A) that contained lens cells are indicated with dotted lines and arrows. Four genes known to be expressed in either lens fiber cells (*cryba1a* (B); *grifin* (C); *mipa* (D)) or epithelial cells (*foxe3* (E)) were used to identify these cell types (F). The intensity of purple coloration in panels B-E reflects gene expression. Panel G shows the developmental age of all lens cells. Note that insets in panels B-G include the 2 and 5 dpf lens epithelial cells. Gene ontology term analysis of the most expressed genes in 1 dpf compared to 2 and 5 dpf lens epithelial cells showed a shift from enrichment in protein translation, ribosome biogenesis and embryonic development at 1 dpf to membrane transport and ATP metabolism at 2 and 5 dpf (Supp Table 1).

The shift in epithelial gene expression between 1 and 2 dpf coincides with the end of lens delamination from the surface ectoderm, transition of the epithelium to a simple cuboidal layer extending towards the posterior pole of the lens, and the initiation of secondary fiber cell formation (Greiling and Clark, 2009). To examine this transition, we identified preferentially expressed genes in the two lens epithelial cell clusters (those with adjusted p-values ≤ 0.05) and submitted them to gene ontology (GO) term analysis using GOrilla (Eden et al., 2009; 2007). Epithelial cells at 1 dpf were enriched in genes related to 10 GO terms associated with protein translation, ribosome biogenesis, and embryonic development (Supplemental table 1). Epithelial cells at 2 and 5 dpf had shifted to 19 GO terms dominated by membrane transport and ATP metabolism (Supplemental table 1). We did not find any noticeable difference in gene expression between fiber cells at 1, 2 and 5 dpf.

### 3.2. Lens crystallin genes show diverse expression patterns

The vertebrate lens includes a diverse group of highly expressed proteins collectively called crystallins that provide the refractive index and transparency required to transmit and focus light onto the retina. Some of these proteins likely also play non-optical roles both within and outside the lens. Our single cell gene expression data provide novel information on the timing and location of crystallin gene expression. These data show how expression has diverged in duplicated crystallin genes, especially in the large number of γ-crystallins.

Zebrafish produce one additional α-crystallin protein compared to mammals because of a duplicated αB-crystallin gene (*cryaba* and *cryabb*) (Smith et al., 2006). There is only one identified zebrafish αA-crystallin gene (*cryaa*) (Runkle et al., 2002). Our scRNA-Seq data showed that zebrafish *cryaa* was expressed almost exclusively in 2 and 5 dpf lens fiber cells (Fig. 2A-B, G). The zebrafish αB-crystallin gene with lens-restricted expression in adults, *cryaba* (Posner et al., 1999), was expressed in a small number of cells located in clusters previously identified (Farnsworth et al., 2019) as lateral line, hindbrain, muscle progenitors and olfactory tissue (Fig. 2C). Expression in each tissue was temporally restricted, with the three hindbrain cells expressing only at 1 dpf, the one olfactory cell at 2 dpf, and the four lateral line (Fig. 2D) and two muscle progenitor cells at 5 dpf. *cryaba* expression was detected in one 5 dpf lens epithelial cell but in no fiber cells. While we report this very restricted *cryaba* expression here, the small number of cells in each tissue with identified transcript may be a technical artifact. The expression of *cryabb* was detected in more cells, with a large cluster identified as hair cells derived from 2 and 5 dpf and a large cluster of otic capsule cells from 5 dpf (Fig. 2E, F). We conclude that *cryaa* expression was lens-focused, and the expression of the two αB-crystallin genes was almost entirely restricted to non-lens tissues during these stages (Fig. 2H).

**Figure 2.**
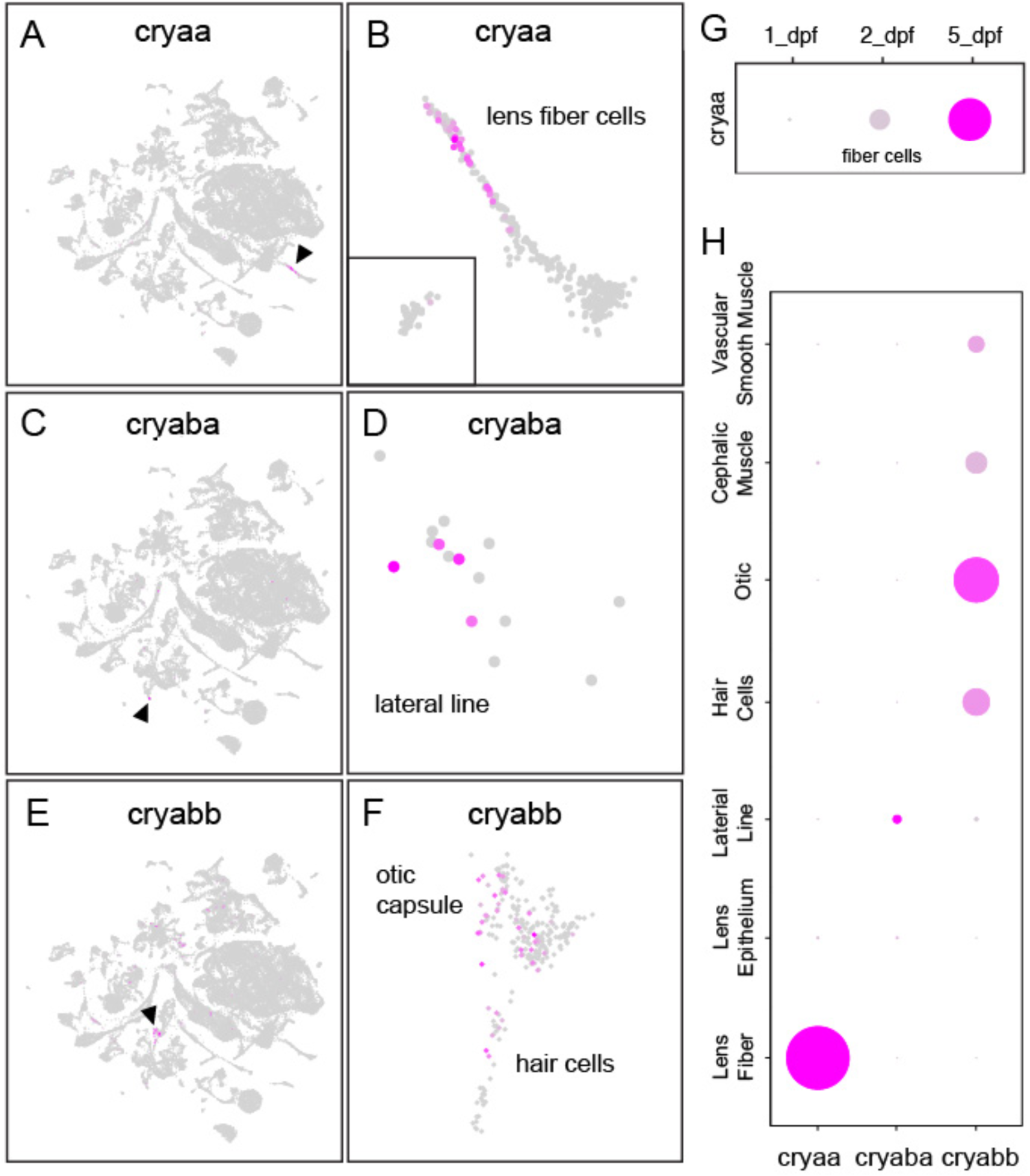
Lens fiber cells upregulated expression of αA-crystallin starting at 2 dpf, but αB-crystallin expression in the lens was almost non-existent through 5 dpf. mRNA for *cryaa* was found almost exclusively in the lens (A), and primarily in fiber cells (B). Expression of *cryaba* was detected primarily in lateral line cells (arrowhead in C and detailed view in D). Expression of *cryabb* was detected at relatively low levels in diverse tissues including otic capsule and hair cells (arrowhead in E and details in F). Temporal expression dynamics of *cryaa* expression in lens fiber cells through 5 dpf (G). Expression of α-crystallin genes in tissues with significant representation (H). For dot plots, the size of each dot reflects the percentage of cells in that tissue expressing the gene while intensity of magenta coloration reflects average gene expression across the tissue cluster.

Little is known about the specific cell types that express zebrafish β-crystallins. We detected expression for 12 of the 13 known zebrafish β-crystallins in our single cell RNA-Seq data, along with four members of the βγ-superfamily (Table 1). Our single cell data match well with publicly available zebrafish developmental whole-embryo bulk-RNA-Seq data (EBI). For example, the EBI bulk-RNA-seq data indicate initial expression for all 13 β-crystallins between 4.3 and 48 hpf, all within the 5 dpf timespan for our single cell data. The one βγ-crystallin that was not found expressed in whole embryos (*crybg2*) was also not identified in our single-cell atlas. Expression of the β-crystallins fit three general patterns: a) expression restricted to fiber cells (multiple βA- and βB-crystallins: Fig. 3A), b) expression primarily in fiber cells but with some expression in epithelium (multiple βA- and βB-crystallins: Fig. 3B) and c) expression outside the lens (for example, *crybb3* was expressed in cell clusters associated with pharyngeal and olfactory epithelium: Fig. 3C). The EBI bulk-RNA-Seq showed that *crybb2* was the latest expressed β-crystallin with mRNA not detected until 48 hpf (EBI). Our single-cell atlas showed *crybb2* expression in several cranial ganglion cells and several lens fiber cells, all of which were 2 and 5 dpf (Fig. 3D). The non-lens crystallin paralogs *crybg1a* and *crybg1b* had very similar expression in periderm, gills, pharyngeal epithelium, otic capsule and hair cells (Fig. 3E-F). One noticeable difference between the two paralogs was greater expression of *crybg1a* in endothelia and of *crybg1b* in notochord cells. Our single cell atlas found *crybgx* expression exclusively in lens fiber cells starting at 2 dpf, and no *crybg2* expression through 5 dpf (Table 1), similar to the whole embryo RNA-Seq data (EBI).

**Table 1.**
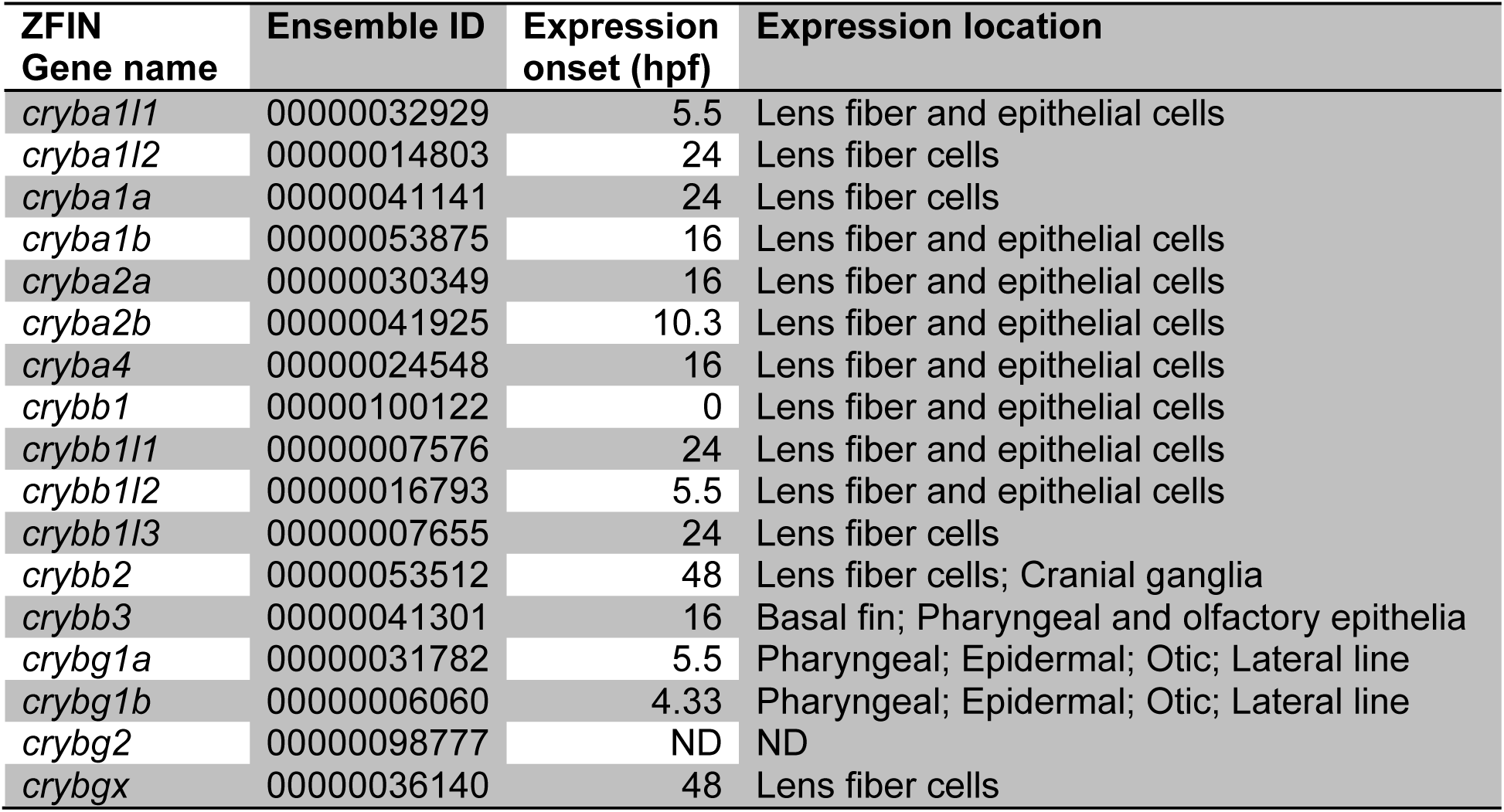
Expression of β-crystallin genes. Age of onset in hours post fertilization (hpf) determined using a publically available whole embryo RNA-Seq dataset (EBI). Location of expression is determined from our single cell RNA-Seq data. Gene expression not detected in either dataset are indicated with ND.

| ZFIN<br>Gene name | Ensemble ID | Expression<br>onset (hpf) | Expression location |
| --- | --- | --- | --- |
| *cryba1l1* | 00000032929 | 5.5 | Lens fiber and epithelial cells |
| *cryba1l2* | 00000014803 | 24 | Lens fiber cells |
| *cryba1a* | 00000041141 | 24 | Lens fiber cells |
| *cryba1b* | 00000053875 | 16 | Lens fiber and epithelial cells |
| *cryba2a* | 00000030349 | 16 | Lens fiber and epithelial cells |
| *cryba2b* | 00000041925 | 10.3 | Lens fiber and epithelial cells |
| *cryba4* | 00000024548 | 16 | Lens fiber and epithelial cells |
| *crybb1* | 00000100122 | 0 | Lens fiber and epithelial cells |
| *crybb1l1* | 00000007576 | 24 | Lens fiber and epithelial cells |
| *crybb1l2* | 00000016793 | 5.5 | Lens fiber and epithelial cells |
| *crybb1l3* | 00000007655 | 24 | Lens fiber cells |
| *crybb2* | 00000053512 | 48 | Lens fiber cells; Cranial ganglia |
| *crybb3* | 00000041301 | 16 | Basal fin; Pharyngeal and olfactory epithelia |
| *crybg1a* | 00000031782 | 5.5 | Pharyngeal; Epidermal; Otic; Lateral line |
| *crybg1b* | 00000006060 | 4.33 | Pharyngeal; Epidermal; Otic; Lateral line |
| *crybg2* | 00000098777 | ND | ND |
| *crybgx* | 00000036140 | 48 | Lens fiber cells |

**Figure 3.**
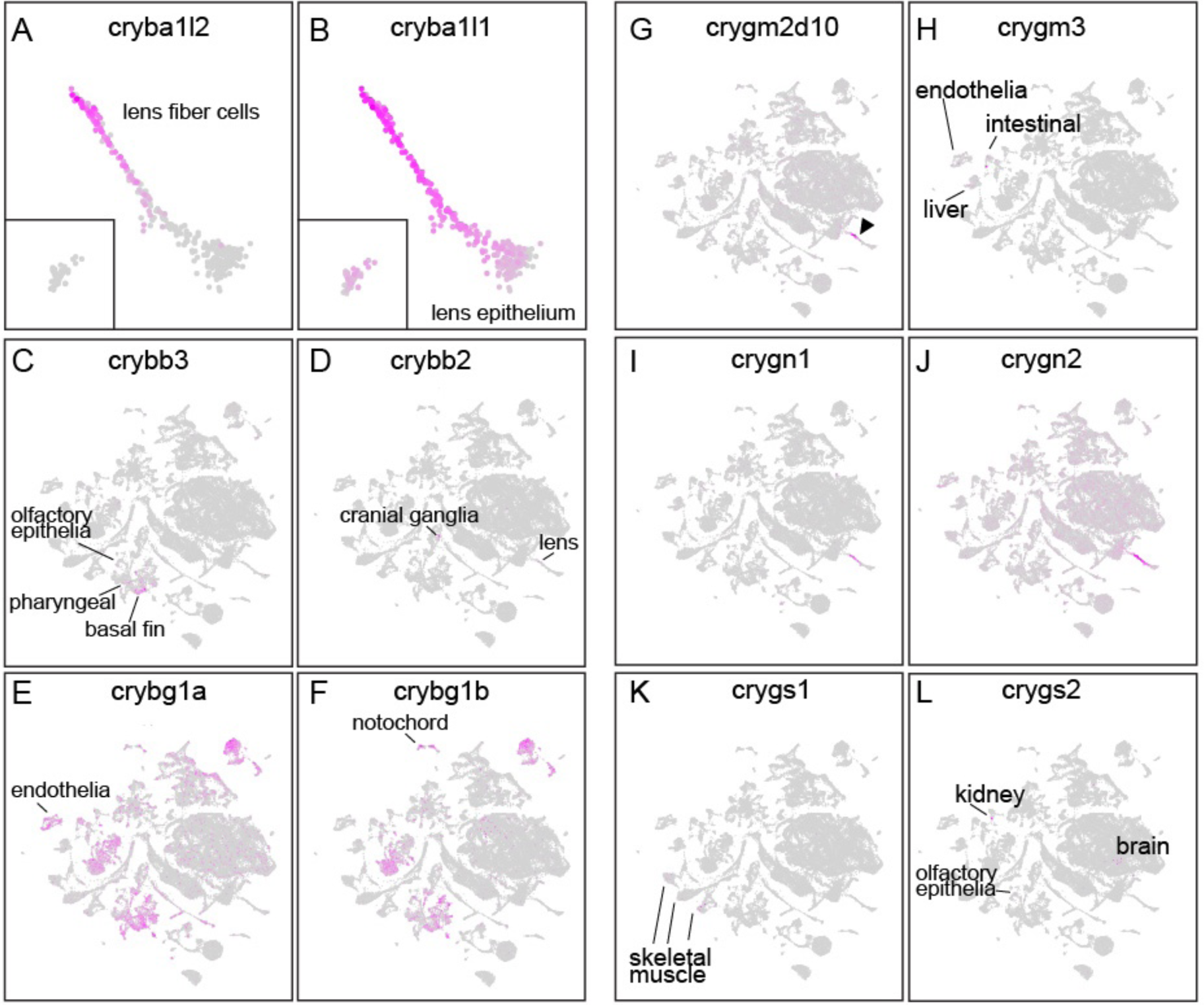
Spatial expression of the β-crystallin and diverse gamma crystallin gene families. Many, like *cryba1l2* were confined to lens fiber cells (A). Others like *cryba1l1* were most strongly expressed in lens fiber cells but also extended into lens epithelium (B). *crybb3* was unique among β-crystallin genes in lacking lens expression and instead being found in olfactory epithelium, pharyngeal and basal fin cells (C). *crybb2* was found in both lens fiber and cranial ganglion cells (D). The paralogs *crybg1a* and *crybg1b* had mostly overlapping expression, although the former was more abundant in endothelia (E) while the latter was abundant in notochord (F). Almost all γM-crystallin genes with detected expression were restricted to lens fiber cells, as shown for *crygm2d10* (G, arrowhead). The only γM-crystallin gene with extralenticular expression was *crygm3*, which was expressed in endothelial, intestinal epithelia and liver cells (H). Both γN-crystallin genes were expressed primarily in lens fiber cells with some lesser expression in lens epithelial cells (I and J). Both were expressed in many other tissues at weaker levels across the embryos, although this expression was stronger for *crygn2*. Only two γS-crystallin genes were expressed in our dataset. *crygs1* was expressed in a number of skeletal muscle cells (K) while *crygs2* was expressed in kidney, brain and olfactory epithelia (L). Neither γS-crystallin gene was expressed in lens.

The γ-crystallins are the largest family of zebrafish lens crystallins due to large gene duplications in the aquatic-specific γM-crystallin group, along with additional duplications in the γN and γS groups. The large number of methionine rich γM-crystallins (34 annotated in zebrafish) has been hypothesized to maintain lens transparency when higher protein densities are required for light refraction in aquatic environments (Kiss and C.-H. C. Cheng, 2008). Zebrafish lack a fourth class found in mammals, the γA-Fs (Wistow et al., 2005). The EBI bulk-RNA-Seq data indicated that unlike a number of β-crystallins that are expressed prior to 20 hpf, the first γM-crystallins are not expressed until 24 hpf (Table 2; EBI). Eleven γM-crystallins were not detected in the EBI bulk-RNA-Seq in whole embryos through 5 dpf (Table 2; EBI). These γM-crystallins whose expression was lacking from the EBI bulk-RNA-Seq data were generally also not detected in our single-cell atlas, indicating concordance with our single-cell atlas. Unlike the expansion of gene expression for several β-crystallin genes into the lens epithelium and outside of the lens, γM-crystallin gene expression was almost entirely restricted to lens fiber cells (Fig. 3G). The only exception was *crygm3*, which showed expression in some endothelial, intestinal epithelia, and liver cells (Fig. 3H).

**Table 2.**
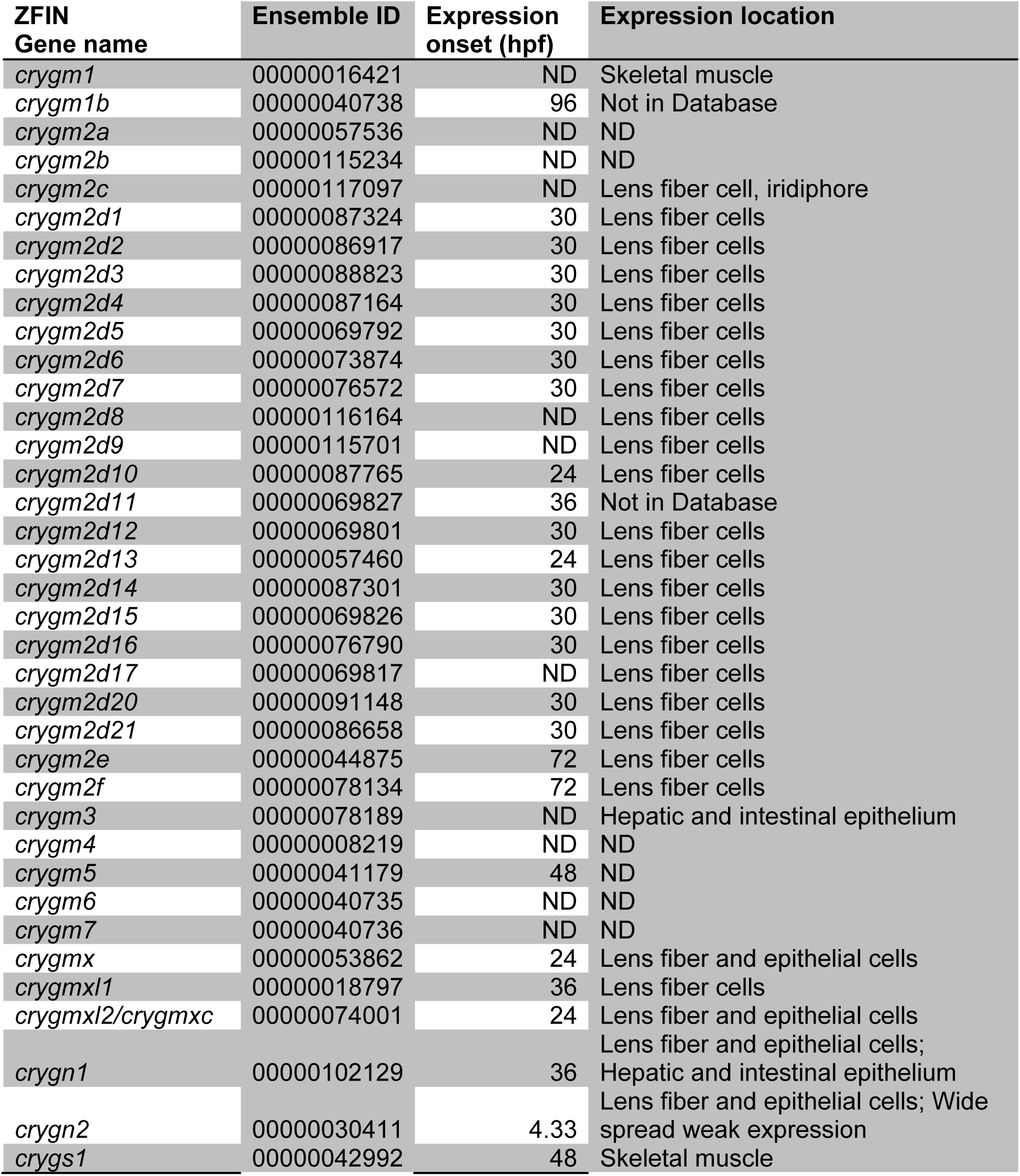
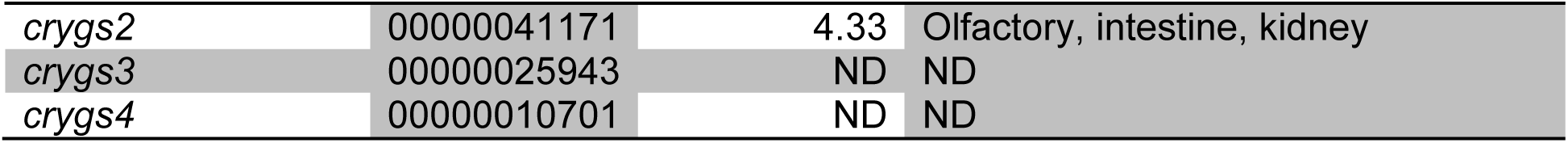
Expression of γ-crystallin genes. Age of onset in hours post fertilization (hpf) determined using a publically available whole embryo RNA-Seq dataset (EBI). Location of expression is determined from our single cell RNA-Seq data. Gene expression not detected in either dataset are indicated with ND.

Zebrafish express two γN-crystallin genes compared to the single mammalian copy, with *crygn2* expressed earlier in development than *crygn1* (EBI) and its product more common in insoluble protein fractions compared to *crygn1*, possibly due to greater interaction with plasma membranes (Posner et al., 2008; Wistow et al., 2005). Our single-cell atlas confirms stronger expression of *crygn2* in the lens through 5 dpf and shows weak expression outside the lens across the entire embryo (Fig. 3I,J). Furthermore, *crygn1* appears more lens specific with some minor expression in cells of the liver and intestine (Fig. 3I). The function of γN-crystallin is not known. It is expressed in mouse but not in human, and in zebrafish the divergence in expression pattern between the two genes suggests a divergence in functions.

The single copy γS-crystallin protein in mammals is expressed as four proteins in zebrafish. The EBI bulk-RNA-seq data showed no expression of *crygs3* and *crygs4* through 5 dpf, but found low expression of *crygs1* at 48 and 72 hpf and low transient expression of *crygs2* at only 4.3 hpf (EBI). Our single-cell atlas shows no expression of any γS-crystallin in the lens through 5 dpf, consistent with our earlier proteomic analysis that could not identify these proteins through 10 dpf (Wages et al., 2013). However, our single-cell atlas did identify *crygs1* transcripts in skeletal muscle cells (Fig. 3K) and *crygs2* in kidney, brain and olfactory epithelia (Fig. 3L). γS-crystallins are expressed only outside the lens prior to 5 dpf.

### 3.3. Zebrafish and mammals share a similar set of lens cytoskeletal and membrane proteins

The vertebrate lens contains a unique set of cytoskeletal and integral membrane proteins (FitzGerald, 1990; FitzGerald and Gottlieb, 1989; Gounari et al., 1993; Hess et al., 1996). The zebrafish lens largely includes the same proteins that are well characterized in mammals. The vertebrate lens expresses two unique intermediate filament protein genes, *bfsp1* (*filensin*) and *bfsp2* (*CP49/phakinin*). Both are found as single copy genes in the zebrafish and their transcripts can be detected starting at 19 hpf in whole embryos (Table 3;EBI). In our single-cell atlas, *bfsp1* is expressed in a small number of lens fiber and epithelial cells while *bfsp2* is restricted to a fairly small population of lens fiber cells (Fig. 4A-D; Table 3). We found five spectrin genes in the single-cell atlas, with *sptan1* most highly expressed in the lens and fiber cell preferred (Fig. 4E). In addition, *sptbn1, sptbn2*, and *sptbn5* were also detected in lens while also broadly expressed across the embryo in multiple non-lens tissues, while *sptb* was specific to red blood cells (data not shown). There is one copy of the vimentin gene (*vim*) in zebrafish and it is abundant in the nervous system, but also found in the lens with more abundance in fiber cells (Fig. 4F).

**Table 3.**
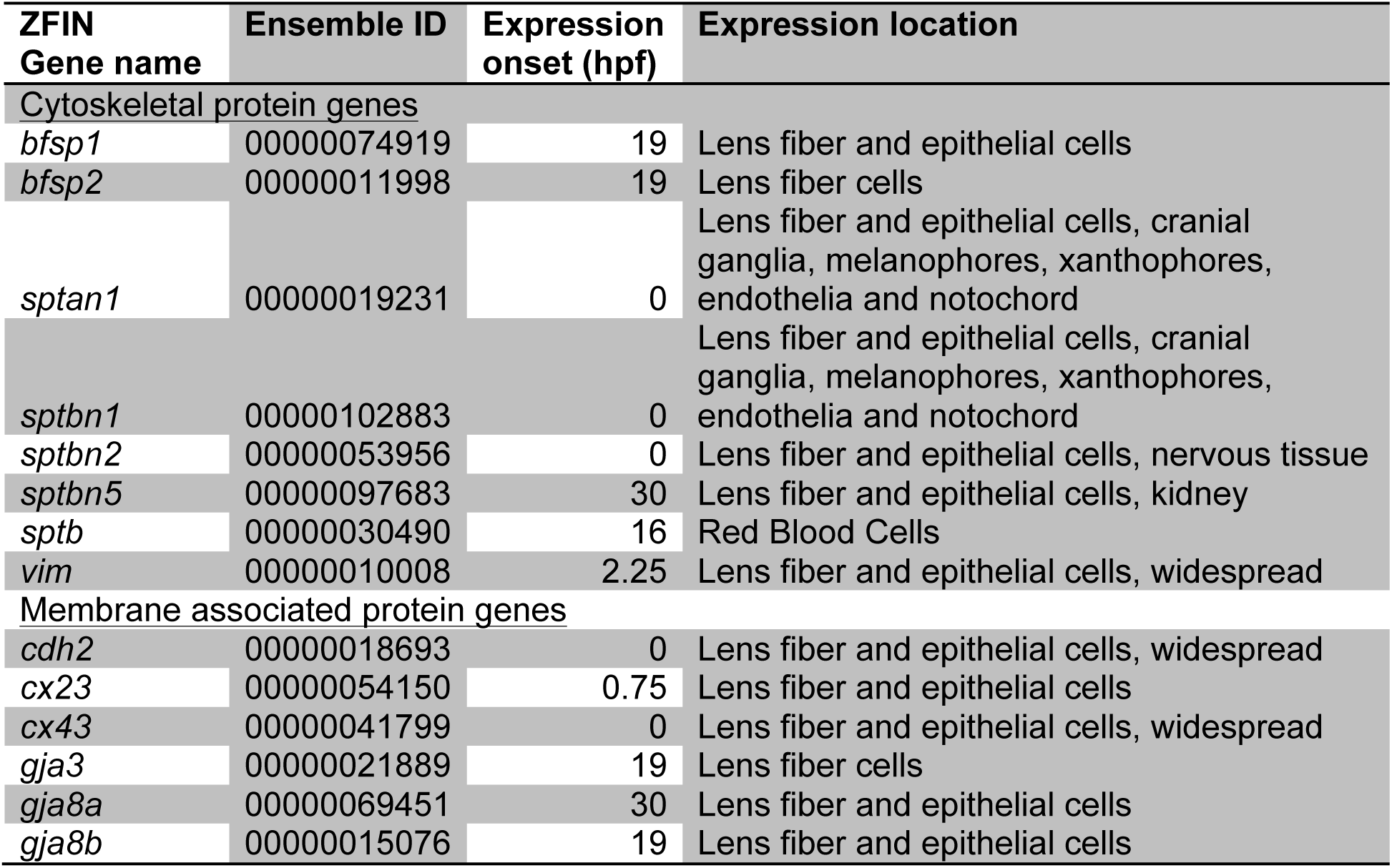
Expression of lens cytoskeletal and membrane protein genes. Age of onset in hours post fertilization (hpf) determined using a publically available whole embryo RNA-Seq dataset (EBI). Location of expression is determined from our single cell RNA-Seq data.

**Figure 4.**
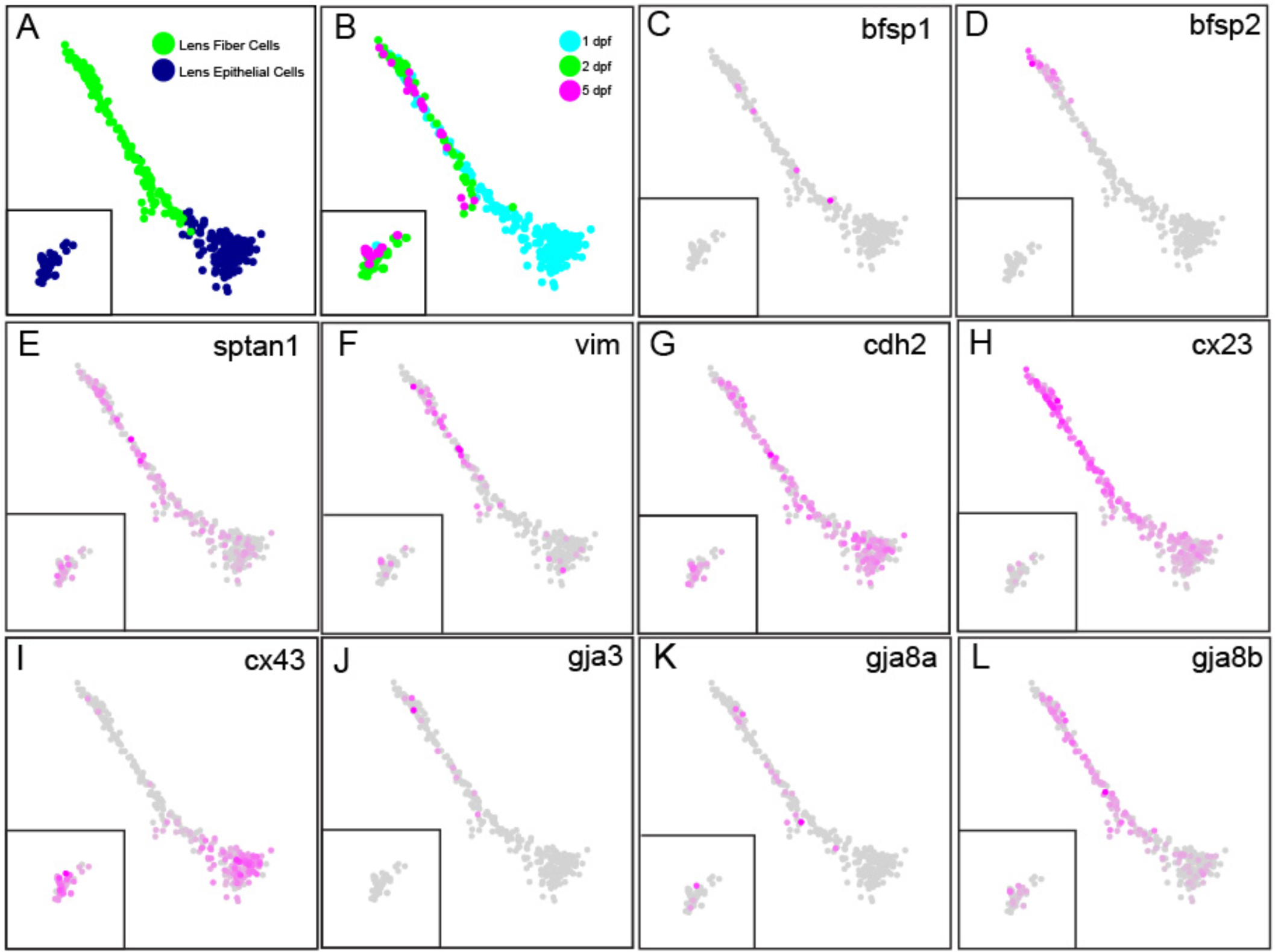
The zebrafish lens expresses many of the same cytoskeletal and membrane proteins as mammals, with some interesting divergence between paralogous pairs. Panels A and B indicate cell type and age as a reference (see Fig. 1). Shown are the expression of two intermediate filament genes unique to the vertebrate lens, *bfsp1* (*filensin*) and *bfsp2* (*CP49/phakinin*) (C and D). Also shown is a spectrin gene (E) and the single vimentin gene (F). Multiple cytoskeletal-associated plasma membrane protein genes are also shown, including one cadherin (G) and two connexin genes (H, I). Three gap junction protein genes are shown, including *gja3* (J) and a paralog pair for *gja8a* and *gja8b* (K, L).

The zebrafish lens for the most part expresses the same contingent of cytoskeletal associated plasma membrane proteins as well studied mammalian models. For example, the zebrafish lens expresses the same single cadherin, *cdh2*, which we found broadly expressed across many cell types, and in lens was found evenly distributed in epithelial and fiber cells (Fig. 4G). Four gap junction/connexin genes were found expressed in the lens. One of them, *cx23* (*gje1*) was only expressed in lens cells, and primarily in fiber cells with some weaker expression in 24 hpf epithelial cells (Fig. 4H). Interestingly, cx23 was not expressed in 2 and 5 dpf epithelial cells. We next analyzed *cx43* expression, which was broadly expressed across the embryo, and in the lens was primarily found in epithelial cells (Fig. 4I). In contrast, *gja3* was expressed almost exclusively in a small number of lens fiber cells (Fig. 4J). The single copy mammalian gene *Gja8* is duplicated in zebrafish with both paralogs almost exclusively expressed in the lens (Fig. 4K, L). The EBI bulk-RNA-seq data showed some temporal differences in expression between *gja8a/*b paralogs: *gja8b* is expressed earlier at 19 hpf compared to 30 hpf for *gja8a* (EBI). Our single-cell atlas also found earlier expression of *gja8b*, in 1, 2 and 5 dpf lens epithelial cells, while *gja8a* was mostly lacking in 1 dpf epithelial cells. Both paralogs were found in lens fiber cells, but *gja8b* was more abundant (Fig. 4K, L).

### 3.4. Spatial and temporal expression dynamics of genes controlling lens development

A number of transcription factors (TFs) involved in vertebrate lens development have been identified, primarily from studies on mammals and birds (see for review Cvekl and Xin Zhang, 2017). While some of these regulatory genes have also been analyzed in zebrafish (for example (Shi et al., 2006; 2005; Swan et al., 2012)), there has been no comprehensive comparison of TF expression patterns between zebrafish and mammalian species. Interestingly, a number of the known mammalian lens development-related regulatory genes exist as paralogs in zebrafish, conserved after the teleost genome duplication. Our single-cell RNA-Seq atlas provides a foundation for examining the expression patterns of known lens-related regulatory genes, possible divergence in the function of paralogs, and the identification of novel regulators.

The best studied lens development regulatory proteins in zebrafish are those that control fiber cell differentiation, such as *foxe3, pitx3* and *hsf4* (Gao et al., 2017; Shi et al., 2006; 2009). Our single cell data showed *foxe3* enriched in lens epithelia (Fig. 5A) while *pitx3* was expressed in both epithelia and fiber cells (Fig. 5B). In the EBI bulk-RNA-Seq data, *hsf4* expression onset is relatively late, beginning at 36 hpf (EBI). Our single-cell atlas identified *hsf4* in a relatively small number of lens fiber cells from 2 or 5 dpf zebrafish (Fig. 5C). The homeobox gene *Prox1* is known to regulate fiber cell differentiation in the mammalian lens (Audette et al., 2016; Wigle et al., 1999). Zebrafish have two paralogs of this gene. Our single cell data show strong *prox1a* expression in lens fiber cells and 24 hpf epithelial cells (Fig. 5D), but almost no lens *prox1b* expression. No study has examined the effects of *prox1a* loss on lens development, although it was shown to play a role in the developmental of lateral line hair cells (Pistocchi et al., 2009). A final TF shown to directly regulate fiber cell differentiation and lens crystallin expression in mammals is *Sox1* (Nishiguchi et al., 1998). A mouse *Sox1* knockout misexpressed *Pax6* during lens development, suggesting a genetic interaction between these transcription factors in the lens (Donner et al., 2007). Zebrafish contain *sox1a/b* paralogs and while their role in neurogenesis has been investigated (Gerber et al., 2019), nothing is known about their role in lens development. Our single-cell atlas showed *sox1a/b* paralogs to have divergent temporal expression in lens epithelial cells, with *sox1a* restricted mostly to 1 dpf epithelial cells while *sox1b* was expressed in 2 and 5 dpf epithelial cells (Fig. 5E, F). Both paralogs were expressed in lens fiber cells.

**Figure 5.**
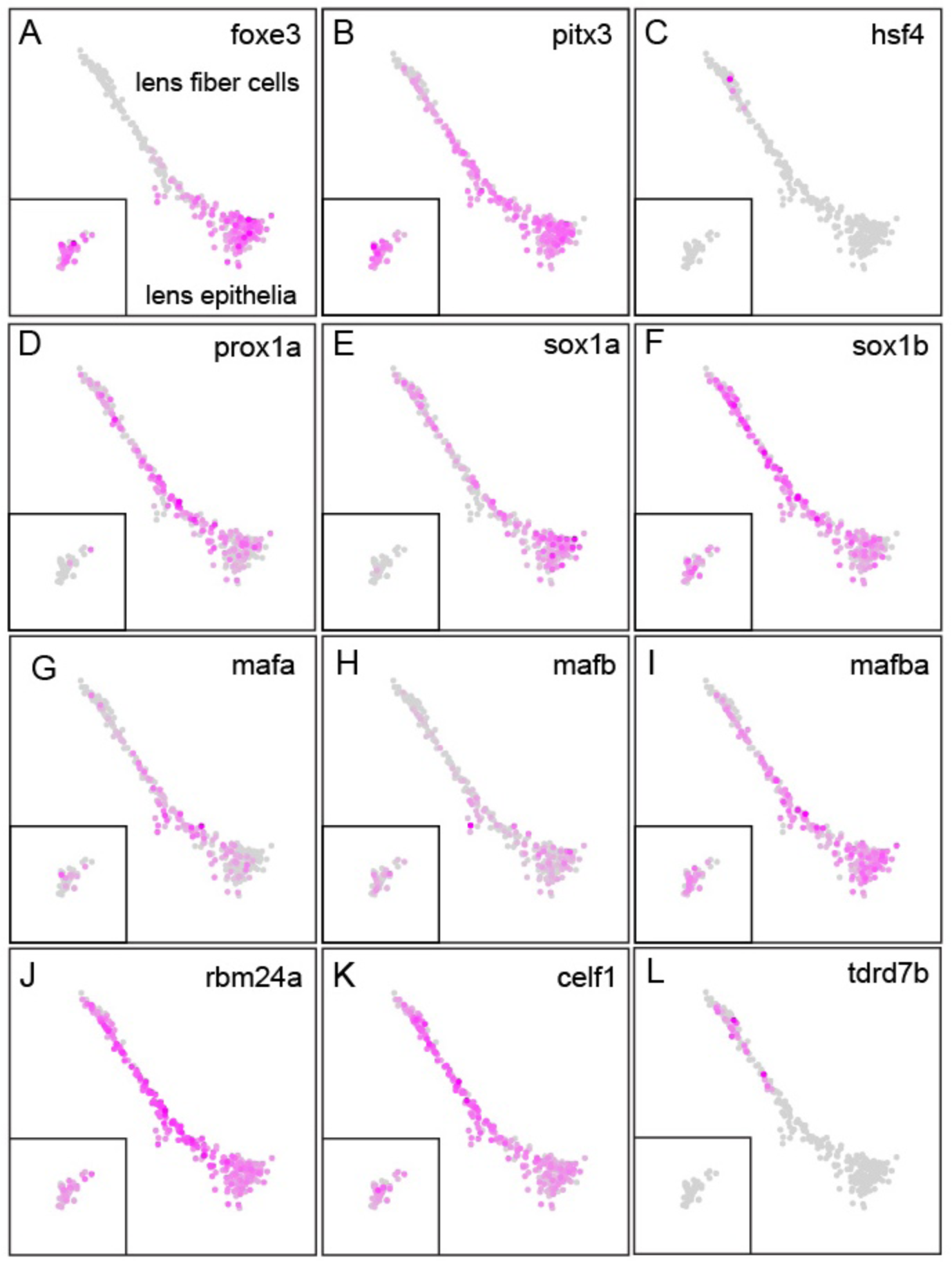
Spatial expression of transcription factors and RNA-binding proteins known to regulate lens development. Expression for the three transcription factors *foxe3* (A), *pitx3* (B) and *hsf4* (C) is shown in the identified lens cell clusters. Inset contains two and five dpf epithelial cells as identified in Figure 1G. These three transcription factors have been shown to take part in shared lens development signaling pathways in both mammalian and zebrafish lens. The mammalian gene *Prox1* is duplicated in zebrafish, but only the paralog *prox1a* is expressed in lens (D). Mammalian *Sox1* also occurs as paralogs in zebrafish, and both are expressed in lens, with *sox1a* showing greater expression in 1 dpf epithelial cells (E) while *sox1b* expression appears to extend into 2 and 5 dpf epithelial cells (F: cells within inset). The expression of three large Maf family genes are shown, including the mammalian c-Maf paralogs *mafa* (G) and *mafb* (H) as well as one mammalian MafB paralog *mafba* (I).Three previously characterized RNA-binding protein genes known to regulate lens development were expressed in lens cells: *rbm24a* (J), *celf1* (K) and *tdrd7b* (L).

The large Maf family of transcription factors are well known regulators of lens development (Cui et al., 2004; Kim et al., 1999; Reza and Yasuda, 2004; Takeuchi et al., 2009; Yang et al., 2004). Vertebrates express four classes of large Mafs with mammals expressing one gene for each class (Coolen et al., 2005). The most critical large Maf in mammalian lens development, c-Maf, occurs as two paralogs in zebrafish, *mafa* and *mafb*, which were both expressed in lens in our single cell data (Fig. 5G, H). Another large Maf, MafB, also occurs as paralogs in zebrafish. But in this case only one of the paralogs, *mafba*, is expressed in lens (Fig. 5I). The last two large Mafs, MafA and NRL occur as single orthologs in zebrafish, with *nrl* expressed in lens but *mafaa* only outside the lens (data not shown).

Almost all of the transcription factors categorized by Cvekl and Zhang (Cvekl and Xin Zhang, 2017) as being involved in mammalian *pax6* signaling, lens vesicle separation, lens cell survival, cell proliferation and differentiation were expressed in lens cells in our single-cell atlas (Fig. 6A,B). The only exceptions, *pknox1.1* and *hif1aa*, have paralog pairs that were expressed in lens. These data indicate a broad retention of lens expression of mammalian-related lens development regulators. Our data also showed that most of these transcription factor genes, unlike the lens-focused expression of genes that directly regulate crystallin expression, were expressed in a broad range of extralenticular tissues (Supp fig. 1). The expression of lens development-related transcription factor genes within the lens varied in both intensity and their relative expression within epithelial or fiber cells (Fig. 6). Most were more strongly expressed in epithelial cells, although *sox2* and *prox1a* were notable exceptions (compare Fig. 6A to 6B).

**Figure 6.**
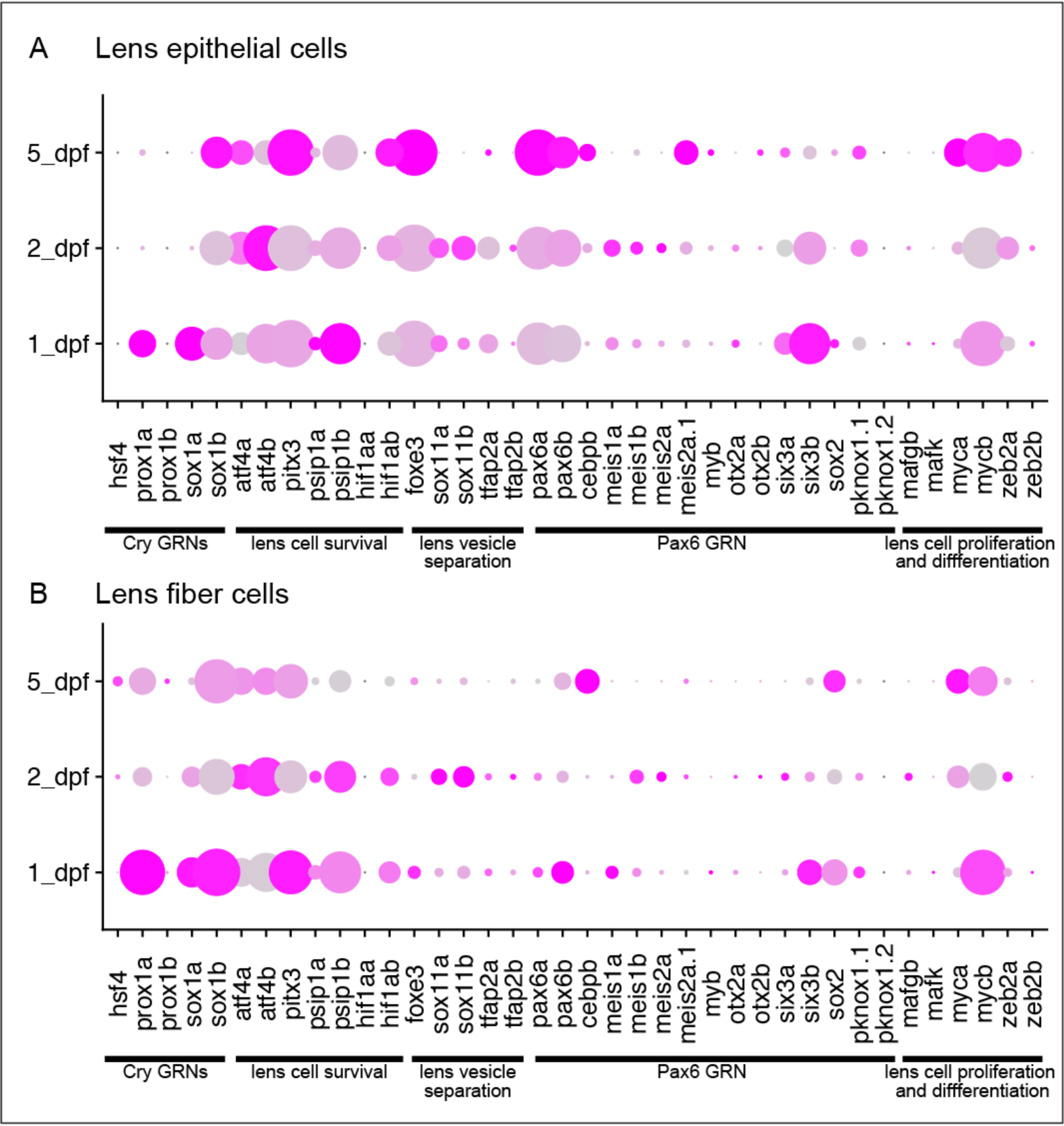
Temporal dynamics of transcription factor expression in zebrafish lens epithelial and fiber cells at 1, 2 and 5 dpf. Expression of lens development regulators and zebrafish paralogs for single copy mammalian genes in lens epithelial (A) and fiber cell (B) clusters. Functional categories for transcription factors are noted below each panel. The size of each dot reflects the percentage of cells in that tissue expressing the gene while intensity of magenta coloration reflects average gene expression across the tissue cluster.

While studies of lens development have focused primarily on the role of transcriptional control of gene expression, recent studies indicate that proteins bind to the untranslated regions of lens mRNAs to modify their expression. Of the over 1600 RNA-binding proteins (RBPs) encoded in the mammalian genome only four have been implicated so far in lens function: Tdrd7 (Lachke et al., 2011), Caprin2 (Dash et al., 2015), Celf1 (Siddam et al., 2018) and Rbm24 (Dash et al., 2020; Grifone et al., 2018). CRISPR-mediated mutation of zebrafish *rbm24a* causes reduced or absent eyes (Grifone et al., 2018). This phenotype may result from the role *rbm24a* plays in regulating lens crystallin mRNA stability (Shao et al., 2020). A recent study showed that morpholino-induced translation blocking of *celf1* causes micropthalmia and visibly cloudy lenses by 4 dpf (Siddam et al., 2018). Our scRNA-Seq data confirm that both genes are strongly expressed in lens (Fig. 5J, K). The functions of *tdrd7a, tdrd7b* and *caprin2* have not been examined in zebrafish. Our data show that *tdrd7b*, but not *tdrd7a*, is expressed in lens (Fig. 5L) and *caprin2* is only expressed in a few lens cells (data not shown).

### 3.5. Identification of possible novel regulators of lens development

Transcriptomic analysis of the mouse lens using RNA-Seq and microarray data has helped identify a number of novel genes involved in lens development (Kakrana et al., 2017; Siddam et al., 2018; Srivastava et al., 2017). These studies hypothesized that genes with lens-specific expression are important for lens function and assessed that function using mutant analysis. This approach has successfully identified a number of genes that regulate lens development, including *Celf1, Tdrd7, Mafg* and *Mafk*. We hypothesized that genes that are important for lens development and function would be strongly enriched in the lens within our scRNA-seq data. To find enriched genes in the developing zebrafish lens, we ranked genes by their preferred expression in either lens epithelia or fiber cells (Supplemental tables 2 and 3). These rank lists include multiple genes already known to be important in lens development, but also identify novel genes.

The top 200 lens fiber cell-preferred genes, ranked by expression relative to all cells outside the cluster, included 12 TF genes and six RBP genes. Of the 12 TF genes in the list, seven are already known to play roles in lens development (*etv4, mafba, mycn, pitx3, prox1a* and *sox1a*/*sox1b*). An eighth, *prox2*, has been knocked out in mice but no lens phenotype was noted (Nishijima and Ohtoshi, 2006). Strong lens expression of *prox2* in zebrafish embryos was also identified by whole mount in situ hybridization (Pistocchi et al., 2008). Morpholino-induced depletion of a ninth gene on our list, *hmx4*, in zebrafish led to a reduced forebrain and narrowed eye field due to disruption in retinoic acid and Sonic hedgehog signaling, but no lens phenotype was examined (Gongal et al., 2011). However, whole mount in situ hybridization analysis did confirm *hmx4* expression in lens (French et al., 2007; Song and Perkins, 2018). The other three TF genes with strong fiber cell preferred expression (*btf3l4, hmx1* and *sox13*) have not been investigated for their role in lens development and provide novel targets for future studies (Fig. 7; Table 4). A SNARE protein gene on the list, *stx3a*, was recently shown to produce a protein isoform in humans that, in epithelial tissue culture, could regulate gene expression through interactions with the transcription factor ETV4 (Giovannone et al., 2018).

**Table 4.** Candidate novel lens-related genes. These genes were identified as having preferred expression in either lens fiber or epithelial cells, or both. The functional category of each gene is listed. Preferred expression was defined as the top 200 expressed gene in lens fiber or epithelial cells. The expression of these genes in lens fiber and epithelial cells is visualized in Figure 7.

| ZFIN<br>Gene name | Ensemble ID | Lens location | Functional Category |
| --- | --- | --- | --- |
| *sox13* | 00000030297 | Fiber | Transcription factor |
| *id3* | 00000054823 | Epithelia | Transcription factor |
| *nr2f6b* | 00000003165 | Epithelia | Transcription factor |
| *btf3l4* | 00000089681 | Fiber and Epithelial | Transcription factor |
| *hmx1* | 00000095651 | Fiber and Epithelial | Transcription factor |
| *endou2* | 00000020122 | Fiber | RNA-binding protein |
| *pabpc4* | 00000059259 | Fiber | RNA-binding protein |
| *serbp1a* | 00000074242 | Epithelia | RNA-binding protein |
| *c1qbp* | 00000039887 | Fiber and Epithelial | RNA-binding protein |
| *endouc* | 00000112234 | Fiber and Epithelial | RNA-binding protein |
| *ssb* | 00000029252 | Fiber and Epithelial | RNA-binding protein |
| *bmp5* | 00000101701 | Epithelia | Signal |
| *slc7a11* | 00000071384 | Fiber and Epithelial | Transport |
| *stx3a* | 00000001880 | Fiber and Epithelial | Protein transport/Transcription factor? |
| *otc* | 00000062147 | Fiber and Epithelial | Metabolism |
| *tkta* | 00000029689 | Fiber and Epithelial | Metabolism |

**Figure 7.**
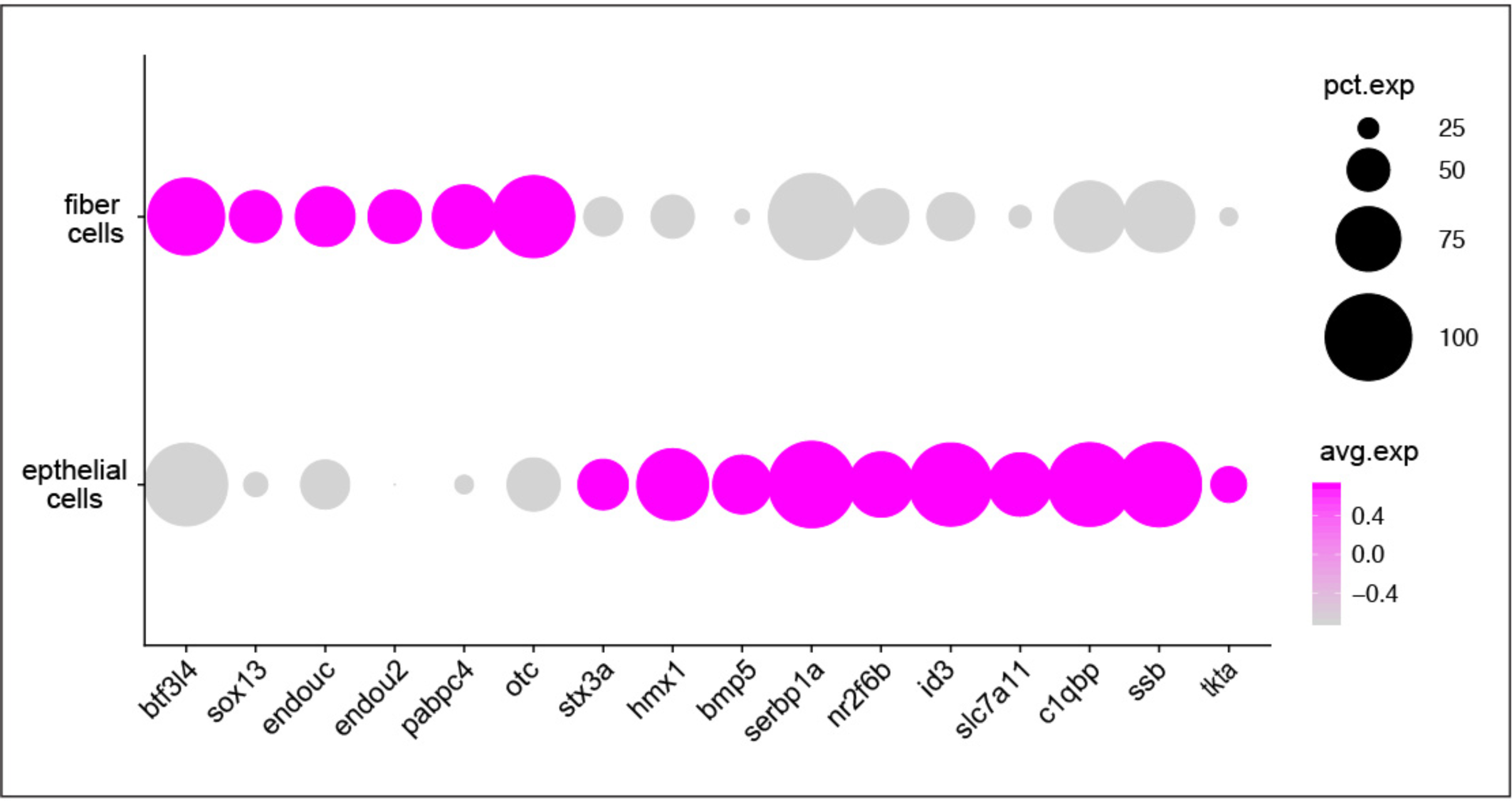
Expression of genes with preferred expression in lens fiber cells and epithelia, but with no currently identified lens function. The size of each dot reflects the percentage of cells in each tissue expressing the gene while intensity of magenta coloration reflects average gene expression across the tissue cluster.

Our list of fiber cell enriched genes also included seven RNA-binding proteins (RBPs), including two that were previously implicated in lens development, *celf1* and *rbm24a* (Fig. 5G and H; Supp Table 2). The other five have no previously described role in lens development or function (Fig. 7). One of them, *pabpc4*, regulates mRNA stability by binding to poly(A) regions, similar to other known lens-related RBPs (D. Liu et al., 2012). Two others (*endou2* and *endouc*) are members of the EndoU ribonuclease family whose members play roles in regulating ER structure, RNA degradation and cell survival (Lee et al., 2017). Available whole mount in-situ data show *endou2* to be expressed strongly in lens by 24 hpf (Thisse and Thisse, 2004), matching our scRNA-Seq data. No role for the EndoU gene family has been previously described for the lens. Another RBP on our list of fiber cell preferred genes, *ssb*, is known to bind to nascent RNA transcripts as well as the poly(A) tail of mRNAs (Vinayak et al., 2018). A final fiber cell preferred RBP, C1qpb, binds mRNA primarily in mitochondria and is linked to several types of cancer and regulated cancer cell chemotaxis and metastasis (Xiaofang Zhang et al., 2013).

A list of the top 200 lens epithelial cluster enriched genes included a large number related to transcriptional machinery, ribosome function and protein synthesis (Supp table 3). There were also multiple genes involved in ATP synthesis and amino acid transport. One amino acid transporter gene, *slc7a11*, showed highly specific expression in lens epithelium and a transketelose (*tkta*) was highly expressed in 2 and 5 dpf epithelial cells (Fig. 7; Table 4). A gene involved in arginine biosynthesis, *otc*, was also highly lens specific and found in both epithelial and fiber cells (Fig. 7; Table 4). We identified five crystallin genes that were enriched in the putative lens epithelial cell cluster, including β-crystallins as well as *crygmx*. Sixteen transcription factor (TF) genes were enriched in lens epithelial cell clusters. Nine of them (*btf3l4, hmx1, hmx4, mafba, pitx3, prox2, sox1a, sox1b, stx3a*) were also enriched in fiber cells (Supp tables 2 and 3). Nine were previously implicated in lens development (*foxe3, mafba, mycb, pax6a, pax6b, pitx3, six3b, sox1a, sox1b*), further showing how our single cell data can highlight functionally important genes. Two additional TF genes (*id3, nr2f6b*) showed lens epithelial-preferred expression but have no previously described lens function (Fig. 7; Table 4). There were six RNA binding proteins (RBPs) in the top 200 epithelial gene list, including five that we also found to be enriched in fiber cells (*c1qbp, celf1, endouc, rbm24a, ssb*). The sixth RBP gene *serbp1a*, was expressed in both epithelial and fiber cells, as well as many tissues outside of the lens and has no known lens function. Lastly, our list of lens epithelial-preferred genes included *bmp5*. While BMP signaling is known to be involved in lens development (Belecky-Adams et al., 2002; Cvekl and Duncan, 2007) *bmp5* specific function has not previously been identified in the lens.

## 4. Discussion

The data presented here are the first single cell level analysis of lens-related gene expression in any species. Lens cells clustered by UMAP non-linear dimensional reduction were identifiable as either fiber or epithelial cells based on the expression of previously identified marker genes. Cell clustering indicated a global shift in gene expression of epithelial cells between 1 and 2 dpf coinciding with secondary fiber cell differentiation. There was no noticeable change in fiber cell global gene expression with age. Our data also provide a detailed description of lens crystallin gene spatiotemporal expression, most significantly clarifying the early expression of the three small heat shock α-crystallin proteins and identifying various patterns of expression in the diverse β- and γ-crystallins. These data provide a new perspective on long-standing questions about lens crystallin evolution and function and possible roles for these proteins outside of the vertebrate lens. Examination of lens cell membrane-associated and cytoskeletal protein genes indicate strong levels of conservation with previously described orthologs from mammalian species, although with some divergence in duplicated genes. Analysis of known lens-related transcription factor genes shows general conservation between mammals and zebrafish, but again with divergence between duplicated paralogs. These single cell expression data identify multiple additional genes with lens preferred expression but with no currently described lens function, including several transcription factor and RNA-binding proteins. In total, our analysis provides a temporospatial atlas of known lens-related genes during early development while also highlighting a number of potential targets for future study.

An important goal of single cell expression analyses is to identify genes with previously undescribed functions in specific tissues. Our lists of lens epithelial and fiber cell preferred genes include at least 16 with no known lens function (Fig. 7; Table 4). Identification of lens preferred expression in the mouse has been a productive approach for identifying new genes with lens related function (Kakrana et al., 2017; Siddam et al., 2018; Srivastava et al., 2017). The majority of the novel genes we identified were for transcription factors and RNA-binding proteins. Follow-up studies on these genes using CRISPR/Cas9 gene targeting could help deepen our understanding of the gene regulatory mechanisms that influence lens differentiation. Identification of novel RNA-binding proteins helping regulate lens development would be particularly impactful as only four such genes have so far been identified for the lens. Several additional genes that could be targeted for study coded for metabolic proteins, possibly leading to new information about lens physiology. The inclusion in our epithelial and fiber cell preferred gene expression lists of multiple genes just recently shown to play a role in lens development, such as the RNA-binding proteins *celf1* and *rbm24a*, supports the hypothesis that the novel genes in our lists are good targets for future gene targeting experiments.

Comparisons between the expression of mammalian and zebrafish lens developmental transcription factors (TFs) are instructive in better understanding their function. Here we summarize three observations about our TF expression data. First, almost every TF known to be involved in mammalian lens development was expressed at some level in zebrafish lens cells. This overall retention of lens expression in the zebrafish orthologs of mammalian lens regulatory genes fits the hypothesis that the mechanics of lens development are conserved within vertebrates. Second, most mammalian lens development-related transcription factors do not show unique or elevated expression in zebrafish lens cells, but instead typically exhibit expression in diverse tissues. For example, almost all of the genes involved in the *pax6* lens regulatory network, including the two zebrafish *pax6* paralogs, show this pattern (Supp fig. 1). Similarly widespread expression was seen in most genes involved in lens cell survival, lens vesicle separation, and cell proliferation/differentiation. While few studies have directly examined whether these genes are involved in zebrafish lens development, it is notable that so many are expressed in lens, while at the same time exhibiting such diverse expression in non-lenticular tissues. Many of the genes known to directly regulate lens crystallin expression (eg. *hsf4, prox1a, sox1a* and *sox1b)* were more specifically expressed in lens cells (Figure 6; Supp Fig 1). The strong levels of lens cell specific expression in this crystallin regulatory group suggests two possible hypotheses: first, crystallin regulation is highly conserved in the vertebrate lens and this evolutionary conservation is reflected in strong gene expression. However, a second possible hypothesis is simply that the large concentrations of crystallin proteins generated by lens cells requires large concentrations of their regulatory transcription factors.

A third and final observation about lens TFs is that the expression patterns of paralogs are often only partially overlapping. For example, while the two *pax6* paralogs have largely overlapping expression in lens, retina, brain and olfactory epithelia, *pax6b* showed unique expression in pancreatic cells (Supp Fig 1). Furthermore, *six3a* and *six3b* showed overlapping expression in forebrain and olfactory cells, but *six3b* was more highly expressed in lens and retina (Supp fig. 1). Most interesting for the study of lens development is the divergent expression between lens epithelial and fiber cells of several paralog pairs. The greater expression of *sox1b* in fiber cells and 2-5 day epithelial cells compared to *sox1a* may reflect subfunctionalization that could be explored through future knockout studies (Fig. 6). A recent study showed co-expression of both zebrafish *sox1* paralogs in brain, lens, otic vesicle and spinal cord by whole embryo in situ hybridization (Gerber et al., 2019). This result is in agreement with the *sox1a/b* expression pattern observed from our single-cell atlas. Relative shifts in expression were also found in the paralog pair *myca*/*mycb*, the orthologs to mammalian C-Myc. For example, while *myca* expression is slightly higher in fiber cells its paralog *mycb* shows a shift to epithelial expression (Fig 6). While they are expressed at lower levels overall, there is similar lens cell expression divergence in the paralogs for *tfap2a/b, otx2a/b* and *meis1a/b*. We do not know if there are any functional consequences to these shifts in expression. In addition, the temporal dynamics of transcription factor expression within lens epithelial and fiber cells found in our single-cell atlas are striking. For example, *pax6a* and *pax6b* show increasing expression over time (1-5 dpf) in lens epithelial cells, but not in fiber cells (compare Fig. 6A to 6B).

The developmental time points we examined surround the key period when secondary lens fiber cells differentiate between 1 and 2 dpf and the epithelium becomes a simple cuboidal layer. During this period we found a shift in lens epithelial cell gene expression from intense protein production to membrane transport and ATP generation (Supp Table 1). This shift may reflect the epithelium’s more terminal role in metabolically sustaining the growing mass of denucleated fiber cells. We do not believe that this early developmental physiological shift has been identified by previous studies. It will be interesting to see if this change in epithelial cell biology is conserved in other vertebrate species as additional single cell analyses are published. Lens fiber cells undergo large morphological changes during the first five days of zebrafish development as secondary fiber cells are added between 1 and 2 dpf and fiber cell denucleation is completed around 3 dpf. However, we did not observe distinct clustering between fiber cells at the three developmental ages examined in our study. Since new fiber cells continue to be produced during lens growth at the stages analyzed in this study, each larva, and thus each sample, may contain both newly born and more differentiated fiber cells that could be transcriptionally similar. It is also possible that strong attachments between maturing fiber cells may bias against their ability to be efficiently profiled using our dissociation methods and scRNA-seq library preparation. In total these data suggest that epithelial cells undergo a significant shift in activity during the first 5 days of zebrafish lens development that is not paralleled in the fiber cells.

A substantial literature on the structure, function and expression of lens crystallin proteins has provided a rich understanding of their roles in lens development, function and disease (Andley, 2007). We believe that our single cell expression dataset provides a valuable, novel perspective on lens crystallins. The most studied lens proteins have been the α-crystallins due to their hypothesized role in maintaining lens transparency and evolutionary relationship with small heat shock proteins (Horwitz, 1992). Our single cell data mirror previous RT-qPCR and RNA-Seq data that indicated an increase in *cryaa* expression starting at 2 dpf (Posner et al., 2017). *Cryaa* expression was detected at an earlier stage in embryonic mouse lens using *in situ* hybridization, where transcripts were found in the lens cup by E10 (Robinson and Overbeek, 1996). This stage is equivalent to approximately 20 hpf in the zebrafish. While we have not captured this stage in our single cell data, publically available whole embryo RNA-Seq data do not show *cryaa* expression at this stage either, suggesting that the zebrafish may delay *cryaa* expression relative to mouse. Later in development mouse *Cryaa* expression becomes most prevalent in fiber cells (Robinson and Overbeek, 1996), similar to what we found in our zebrafish single cell data.

There are conflicting data in the literature about the early expression pattern of the two zebrafish αB-crystallins (see (Posner et al., 2017) for discussion). Briefly, RT-PCR and qPCR of whole embryos did not detect expression of *cryabb* through 78 hpf and 5 dpf, respectively, while qPCR did detect *cryaba* starting at 12 hpf (Elicker and Hutson, 2007; Mao and Shelden, 2006). We previously showed a similar pattern by qPCR, with *cryaba* expression increasing between 4 and 5 dpf but minimal expression of *cryabb* (Posner et al., 2017). Another recent study detected both aB-crystallin genes starting at 24 hpf (Zou et al., 2015). Our single cell RNA-Seq data help address these discrepancies by identifying the embryonic cell types expressing each gene. No *cryabb* mRNA was detected in lens cells through 5 dpf and only one lens cell, a 5 dpf epithelial cell, expressed *cryaba*. Our data suggest that both αB-crystallin mRNAs are essentially non-existent in the lens through 5 days of development. However, both genes were expressed at varying levels outside the lens. What functional role, if any, these proteins play in those cells is not known. In adult zebrafish *cryaba* becomes mostly lens-specific while *cryabb* is expressed broadly, similar to the mammalian ortholog (Smith et al., 2006). The timing of these ontogenetic shifts in expression, and when exactly both αB-crystallin genes are expressed in lens, will require examination of older developmental stages. It is interesting that in embryos, as in adults, *cryabb* has the most diverse tissue expression of the three a-crystallin genes. Neither zebrafish αB-crystallin gene shares the same expression pattern as mouse αB-crystallin, which was detected in the lens placode of mouse embryos and became preferentially expressed in the lens epithelium as primary lens fiber cells differentiated (Robinson and Overbeek, 1996), a pattern quite different from that of the zebrafish.

It is possible that αB-crystallin proteins may be accumulating in lens cells without detectable levels of mRNA, or that we are simply failing to detect their mRNAs (a false negative), but considering that we were able to detect mRNA for a number of other lens crystallins at these same ages this interpretation seems unlikely. A previous mass spectrometry examination of zebrafish lenses found neither αB-crystallin in the lens through three weeks of age, but abundant protein at 6 months of age (Greiling et al., 2009). While future proteomic examination of embryonic zebrafish lenses would provide additional data regarding the timing of αB-crystallin accumulation in the lens, it appears that these proteins are not present during early development. We previously showed that the proximate promoter regions for the two zebrafish αB-crystallins did not efficiently drive GFP expression in the embryonic lens, mirroring our new scRNA-seq data (Posner et al., 2017). However, that study also showed that both zebrafish αB-crystallin promoters drove GFP expression in skeletal muscle and notochord in embryos at 2 dpf, a result that is not reflected in our single cell data. It is possible that more distal enhancer regions alter the expression of these genes, providing a caution when interpreting analysis of isolated promoter regions.

Much less is known about the function of β- and γ-crystallins in the vertebrate lens. Most work on this group has focused on the impact of mutations, post-translational modifications and protein aging in cataract development (Lampi et al., 2014), although some studies have examined the roles of β-crystallins in retinal astrocytes and in phagocytosis within the retinal pigmented epithelium (Sinha et al., 2008; 2012; Zigler et al., 2011). Zebrafish have a similar contingent of β-crystallins compared to mammals, but with added genes due to duplications (Wistow et al., 2005). The contingent of zebrafish γ-crystallins is quite different from mammals due to the presence of the aquatic lens γM-crystallins and loss of the ϒA-F-crystallins. No study has presented comprehensive data on the spatial expression of β- and γ-crystallins in any species.

Mammalian lenses express eight β-crystallin proteins using seven genes (Wistow, 2012). Zebrafish have expanded this family to 13 genes, seven βA- and six βB-crystallin genes. The largest expansion is of mammalian βA1-crystallin. While this one gene produces two different proteins in mammals, zebrafish have four βA1 genes encoding four known proteins. The two paralogs expressed earliest in development (*cryba1l1* and *cryba1b*) also showed more extensive lens expression than the other two f3A1-crystallins, extending from fiber cells into epithelia. Future work will need to explore whether these differences in expression are connected to variation in function. The largely overlapping expression of the two zebrafish βA2 paralogs does not suggest functional divergence, but this question would also need to be examined experimentally. The single βB1 mammalian crystallin is found as four genes in zebrafish. Onset of expression as shown in whole embryo RNA-Seq data is quite divergent between the four genes, which may suggest unique roles of these paralogs. The spatial range of expression from fiber cells into epithelial cells also differed. The single βB2 gene showed limited expression in lens fiber cells as well as expression in a few cells identified as cranial ganglia. The only βA- or βB-crystallin with significant expression outside the lens was βB3-crystallin. This observation is interesting considering that mammalian *Crybb3* is also unique among f3-crystallins in being expressed transiently within the nuclei of differentiating lens fiber cells (Limi et al., 2019).

No studies have examined the role of the fish specific β-crystallin *crybgx*, and its very limited expression in our dataset (only found in two lens fiber cells) does not suggest a vital role in lens development. Zebrafish contain three genes for the non-lenticular members of the β-crystallin family. The encoded proteins are often not referred to as crystallins since they are not produced in the lens, and in humans are known as the AIM1 and AIM1-like proteins. The two zebrafish paralogs for human *AIM1, crybg1a* and *crybg1b*, showed largely overlapping expression outside the lens, again suggesting conservation of function between paralogs.

Despite the large number of zebrafish γM-crystallin genes, most of them showed remarkably similar expression, with onset around 30 hpf when secondary fiber cells begin to differentiate. These data showing late and lens specific use support the hypothesis that the role of γM-crystallins is restricted to a building material allowing for close packing of proteins in a high density aquatic lens. According to this hypothesis, diversity in protein shape provides for the polydispersity that helps prevent protein aggregation and opacity at high density. While α-crystallins achieve this polydispersity by forming a range of aggregate sizes (Aquilina et al., 2005) γ-crystallins form uniform dimers, so multiple proteins may be needed to achieve this same effect in the aquatic vertebrate lens. While the diversity of γM-crystallins is not know from the sarcopterygian fishes that are more closely related to tetrapods, it is possible that this entire class of ϒ-crystallins was lost during the transition of vertebrates to land.

Possible roles played by the other two classes of ϒ-crystallins remain unclear and a potentially interesting open question. The lack of mRNA for any of the four zebrafish γS-crystallins in the lens through 5 dpf matches previous proteomics studies (Greiling et al., 2009; Wages et al., 2013). Together these results suggest that ϒS-crystallins play no role in lens development, but do become abundant members of the adult lens proteome (Posner et al., 2008). While their role in the lens is yet to be determined, the early expression of *crygs1* in skeletal muscle and *crygs2* in olfactory and kidney tissue may indicate non-ocular developmental roles. Further study of these two genes may help us understand the roles of the single copy γS-crystallin in mammals. Likewise, the two zebrafish paralogs of the single copy mammalian γN-crystallin have diverged in timing of expression onset and extent of lens expression. The function of these proteins in any species is largely unknown, making zebrafish an excellent model for exploring their biological roles. It is worth noting that many of the βA-, βB- and γM-crystallins showed very weak expression in a wide range of embryonic tissues. This pattern was not seen in α-crystallins, suggesting that it is specific to these crystallin classes. Whether this weak background expression reflects any functional importance in these tissues is not known.

Unlike the large numbers of paralog pairs in the crystallins and lens regulatory genes, zebrafish membrane associated and cytoskeletal proteins occur mostly as one-to-one orthologs with mammals. The only exception is the presence of two paralogs for *gja8*, which did differ in expression and could possibly be useful for studying this protein. The single copy *gja3* (AKA cx48.5) provides an interesting test of how well different experimental approaches agree when analyzing gene expression and function. A previous study identified *gja3* mRNA by RT-PCR in 4 dpf zebrafish lens and heart, while whole-mount in-situ hybridization found mRNA in lens and transiently in otic vesicles, but not in heart (S. Cheng et al., 2004). Our scRNA-Seq data found *gja3* expressed in lens fiber cells, with expression in some cells of the periderm and gills, but not in heart or otic vesicles (Fig. 4J, some data not shown). Interestingly our single cell data and the previous study’s in-situ hybridizations both showed that lens *gja3* expression is restricted to fiber cells. Morpholino-induced reduction of *gja3* expression produced lens abnormalities and cardiovascular defects in 3 dpf embryos (S. Cheng et al., 2004). Further comparisons of data resulting from different gene expression analysis approaches (ie. RT-PCR, RT-qPCR, RNA-Seq, scRNA-Seq, promoter analysis) combined with functional testing of gene function using translation blocking and knockout tools will help to clarify how well gene expression levels predict functional importance in a specific tissue. Throughout our results section we have attempted, when possible, to compare our single cell RNA-Seq data to published data from in situ hybridization qPCR and bulk RNA-Seq experiments to determine the level of concordance between techniques. We generally found that these data match, but recommend that any follow up analyses of gene expression patterns described in our single cell dataset should be confirmed by in situ hybridization analysis. In the case of *gja3* it is interesting that some techniques showed similar expression in some tissues, but not others. The demonstration that reduction in *gja3* expression affects heart development is not predicted by expression patterns resulting from in situ hybridization and single cell RNA-Seq. It is worth noting that single cell RNA-Seq can lead to false negatives if transcripts are not captures, but false positives are unlikely. Therefore, identification of novel genes not previously shown to have lens preferred expression is likely a useful approach for selecting targets for future analysis of lens development and function.

We noticed a number of lens-related genes that shared expression with tissues derived from the preplacodal region. This vertebrate embryonic tissue is a shared developmental origin for the lens and hair cell-containing sensory organs like the inner ear, olfactory epithelium and lateral line (Bailey and Streit, 2005; Grocott et al., 2012). Examination of the molecular signaling that leads to differentiation of these different sensory structures showed that the default gene expression pattern leads to lens development, with the other structures requiring shifts in the production of signaling molecules (Bailey et al., 2006). Study of vertebrate tissue regeneration also shows shared molecular signaling pathways between the lens and inner ear hair cells (Tsonis, 2007; Tsonis et al., 2007). Both αB-crystallins were expressed in extralenticular tissues of preplacodal origin with *cryaba* expressed primarily in one of these tissues, the lateral line, while *cryabb* was found in lateral line, otic capsule, and other hair cells (Fig. 2H). One β-crystallin, *crybb3*, was expressed in olfactory epithelia, two β/γ-crystallins were expressed in otic capsule and lateral line and one γ-crystallin, *crygs2*, was expressed in olfactory epithelium. Interestingly, all of these crystallins with extralenticular preplacodal tissue expression are either non-existent in lens or expressed at lower levels in the lens. This observation suggests that these genes may be pulled away from the lens function of their other crystallin family members by changes in gene expression signaling into other roles in preplacode derived tissues.

Transcription factor genes have also been shown to share expression in placode-derived tissues. Zebrafish *pitx3* was found to be expressed in placode regions that give rise to the lens, olfactory epithelium, anterior pituitary and cranial ganglia, but not the otic and lateral line placodes (Zilinski et al., 2005). Our single cell data mostly agree, showing *pitx3* most highly expressed in lens epithelium, lens fiber cells and hypophysis, but little expression in hair cell tissues (Supp fig. 1). However, we saw minimal expression in olfactory tissue. We did find a number of lens-related TF genes expressed in a cell cluster identified as olfactory epithelium/placode at greater levels than their general widespread expression (e.g. *pax6a, cebpb, meis1a, six3a* and *six3b*; Supp fig. 1). The same set of genes showed shared increased expression in lateral line and hair cells (*sox1a, sox1b, tfap2a, cebpb, meis1a, meis1b, sox2* and *mycb*), while several genes showed shared expression in hypophysis (*pitx3, tfap2a, tfapsb* and *cebpb*). There are opportunities to further explore the connection between signaling molecules leading to diversification of placode-derived tissues.

The work presented here provides an embryo-wide view of the expression of genes linked to lens development and function. We do not know of previous studies in any species that characterize the expression of such diverse families of lens genes at single cell resolution. The data address questions about α-crystallin expression and provide the first systemic analysis of zebrafish β- and γ-crystallin expression. The large number of β- and γ-crystallin paralog genes, and the presence of the unique γM-crystallins in the fish lens, allow for a new view of the evolution of these gene families in the vertebrate lens. Analysis of cytoskeletal, membrane and transcription factor genes supports the hypothesis of strong conservation between mammal and zebrafish lenses, while pointing to areas where gene duplication in zebrafish may provide opportunities to further dissect the roles of these genes. Lastly, the identification of genes with preferred expression in lens, sometimes specific to one cell type, shows promise for identifying novel regulators of lens expression, including new RNA-binding proteins, whose functions in lens development have only recently been recognized.

## Supporting information

Suuplemental Figure 1

## Figure and Table Legends

**Supplemental Figure 1. Expression levels of known regulators of mammalian lens development across all zebrafish cell types**. This dot plot arranges lens development regulators identified in mammalian studies by function as defined in Cvekl and Zhang (2017). Known zebrafish paralogs for single copy mammalian genes are included. Size of each dot reflects expression levels. Cluster numbers can be connected to hypothesized tissue types using supplemental table 4.

**Supplemental Table 1. Gene Ontology (GO) term comparison of 1 dpf and 2-5 dpf lens epithelial cells**. Lists of preferentially expressed genes in each cluster were submitted to GOrilla. Shown are the GO term ID and descriptor, gene count for each GO term, p-value returned by GOrilla and the top genes for each term (up to a total of five). Only GO terms with a p-value ≤ are shown.

**Supplemental Table 2**. Top 200 genes expressed in a cell cluster identified as lens fiber cells ranked by their log fold change in expression compared to cells outside the cluster. Shown along with gene name are adjusted p-values, fraction of cells within and outside the lens cluster with detected expression, the ratio of that expression, the general category of each gene’s function and whether the gene was also in the top-200 expressed lens epithelial genes.

**Supplemental Table 3**. Top 200 genes expressed in a cell cluster identified as lens fiber cells ranked by their log fold change in expression compared to cells outside the cluster. Shown along with gene name are adjusted p-values, fraction of cells within and outside the lens cluster with detected expression, the ratio of that expression, the general category of each gene’s function and whether the gene was also in the top-200 expressed lens epithelial genes.

**Supplemental Table 4**. List of hypothesized cluster identifications by number.

## Acknowledgements

We would like to thank Adil Hussen for help with the analysis of crystallin gene expression and the Busch-Nentwich lab for providing RNA-seq data. Funding: This work was supported by the NIH National Eye Institute [R15 EY13535] to M.P. and by the NIH Office of the Director [R24 OD026591], NIH National Institute of Neurological Disorders and Stroke [R01 NS105758], and the University of Oregon to A.C.M. Funding sources played no role in study design; in the collection, analysis and interpretation of data; in the writing of the report; or in the decision to submit the article for publication.

## Notes

### Competing Interest Statement

The authors have declared no competing interest.

### Summary of Updates

Figure sizes improved for easier viewing

