## Supplementary figures and images for "Single cell transcriptomics of the developing zebrafish lens and identification of putative controllers of lens development"

### Suuplemental Figure 1

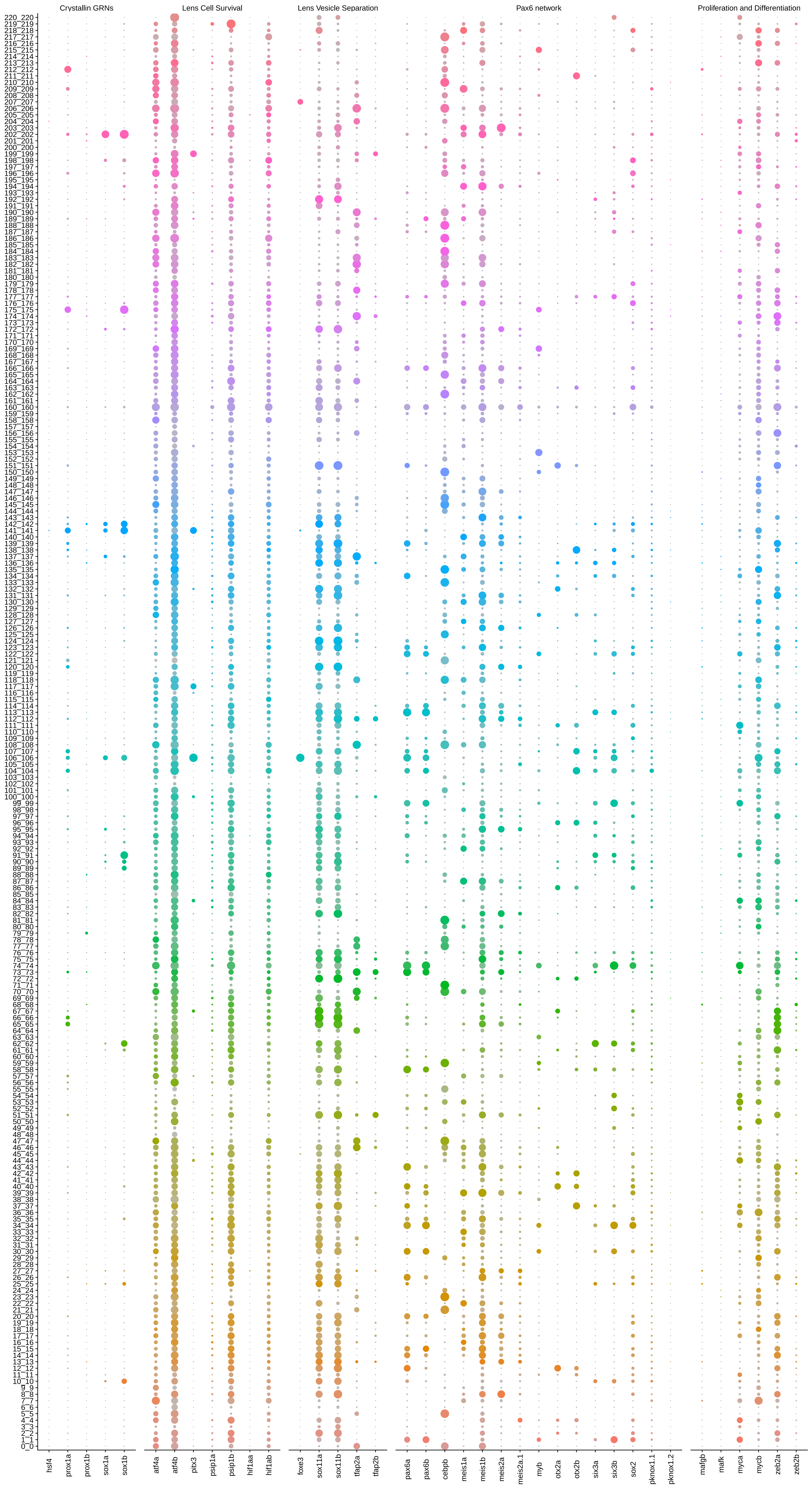
