## Supplemental Table 1 for "Single cell transcriptomics of the developing zebrafish lens and identification of putative controllers of lens development"

**Supplemental Table 1. Gene Ontology (GO) term comparison of 1 dpf and 2-5 dpf lens epithelial cells**. Lists of preferentially expressed genes in each cluster were submitted to GOrilla. Shown are the GO term ID and descriptor, gene count for each GO term, p-value returned by GOrilla and the top genes for each term (up to a total of five). Only GO terms with a p-value ≤ 0.05 are shown.

| GO Term | Count | p-value | Top Genes |
|---|---|---|---|
| | | ***1 day post fertilization lens epithelium*** | |
| **0006412 Translation** | 15 | 2.96E-11 | RSL1D1, EEF2B, EIF3JB, EEF1B2, RPL32 |
| **0002181 Cytoplasmic translation** | 6 | 3.35E-08 | EIF3JB, RPL7, RPLP0, RPL8, EIF4BB |
| **0000470 Maturation of LSU-rRNA** | 4 | 4.46E-05 | RSL1D1, NOP2, RPL7A, NSA2 |
| **0006414 Translational elongation** | 4 | 2.32E-04 | EEF2B, EEF1B2, EEF1DB, EEF1A1L2 |
| **0042254 Ribosome biogenesis** | 4 | 0.0012 | GTPBP4, RPLP0, RPL7A, RSL24D1 |
| **0006364 rRNA processing** | 4 | 0.0022 | NOP2, DKC1, NCL, RPS7 |
| **0043009 Chordate embryonic development** | 4 | 0.0155 | EEF2B, RPLP0, RPS4X, RPS7 |
| **0042274 Ribosomal small subunit biogenesis** | 2 | 0.0288 | SURF6, RPS7 |
| **0000154 rRNA modification** | 2 | 0.0287 | NOP58, NOP56 |
| **0051726 Regulation of cell cycle** | 3 | 0.0291 | RPL7, MDKA, RPS7 |
| | | ***2 and 5 day post fertilization lens epithelium*** | |
| **0015986 ATP synthesis coupled proton transport** | 9 | 1.27E-11 | ATP5E, ATP5B, ATP5C1, ATP5L, ATP5G1 |
| **0015992 Proton transport** | 5 | 8.69E-05 | ATP5B, ATP5C1, ATP5G1, ATP5A1, ATP5J |
| **0006810 Transport** | 22 | 1.04E-04 | MIPB, ATP5J, UQCRB |
| **0006754 ATP biosynthetic process** | 3 | 0.0016 | ATP5B, ATP5C1, ATP5A1 |
| **0006096 Glycolytic process** | 4 | 0.0024 | PFKPA, GAPDHS, PGK1, ENO1A |
| **0015991 ATP hydrolysis coupled proton transport** | 4 | 0.0043 | ATP1A1A.2, ATP5B, ATP5G1, ATP5A1 |
| **0006123 Mitochondrial electron transport, cytochrome c to oxygen** | 3 | 0.0050 | COX8A, COX6A1, COX5AB |
| **0055114 Oxidation-reduction process** | 12 | 0.0075 | GAPDHS, CYP2AD3, CYP26C1, PRDX6, PHGDH |
| **0015671 Oxygen transport** | 3 | 0.0091 | HBBE1.1, HBBE2, HBAE3 |
| **0060142 Regulation of syncytium formation by** | 2 | 0.0152 | SLC3A2A, SLC3A2B |

| | | | |
|---|---|---|---|
| **plasma membrane fusion** | | | |
| **0055014 Atrial cardiac muscle cell development** | 2 | 0.0152 | CYP26C1, CYP26A1 |
| **0003333 Amino acid transmembrane transport** | 3 | 0.0277 | SLC38A2, SLC7A11, SLC7A3A |
| **0006811 Ion transport** | 9 | 0.0292 | SLC38A2, ATP1A1A.2, KCNK2A, ATP5C1, ATP1B1A |
| **0021661 Rhombomere 4 morphogenesis** | 2 | 0.0302 | CYP26C1, CYP26A1 |
| **0042573 Retinoic acid metabolic process** | 2 | 0.0302 | CYP26C1, CYP26A1 |
| **0031175 Neuron projection development** | 3 | 0.0357 | NCAM1A, STMN1A, STMN1B |
| **0006750 Glutathione biosynthetic process** | 2 | 0.0377 | GCLC, GCLM |
| **0055113 Epiboly involved in gastrulation with mouth forming second** | 3 | 0.0409 | APOA2, CFL1, SDC4 |
| **1902600 Hydrogen ion transmembrane transport** | 2 | 0.0450 | COX8A, UQCRB |
